# Comparative Analysis of Relative Ligand Binding Free Energy Simulation Methods: Amber-TI, GROMACS-NETI, OpenMM-FEP, and BLaDE-MSLD

**DOI:** 10.64898/2026.04.22.720125

**Authors:** Hajin Lee, Injung Kim, Sungjun Kim, Minsik Bae, Bomi Jeong, Seonghoon Kim, Sunhwan Jo, Jumin Lee, Wonpil Im

## Abstract

Structure-based drug design has become increasingly important in the pharmaceutical industry for accelerating the discovery of effective drug candidates. In particular, ligand binding free energy serves as a critical metric for predicting drug efficacy during the key stages of hit discovery and lead optimization. Continuous progresses have been made in the prediction of ligand binding free energies, but direct comparisons of different methods using the same force field remain challenging due to their unique implementations into different simulation engines. In this study, we present a direct comparison of four popular methodologies (Amber-TI, GROMACS-NETI, OpenMM-FEP, and BLaDE-MSLD) for calculating relative binding free energies (ΔΔG_bind_) with the same Amber protein and ligand force fields using *MolCube Alchemical Free Energy Simulator* (*MolCube-AFES*), which provides an input generation workflow to support ΔΔG_bind_ calculations of all four methods. We used 80 alchemical transformations (among the JACS benchmark set by *Wang et al.*) and two additional applications to compare the predicted ΔΔG_bind_ from the four methods against experimental measurements. All four methods reproduced experimentally observed trends with most transformations within ±2 kcal/mol from experiments and show broadly comparable accuracy with no statistically significant performance differences across the benchmark dataset. These results demonstrate that *MolCube-AFES* enables controlled, cross platform benchmarking and show that all four different alchemical free energy methods deliver statistically equivalent accuracy, with method selection guided by workflow requirements such as throughput, portability, and perturbation network design rather than expected differences in performances.

## 1. Introduction

Free energy calculations are essential tools in molecular simulations, providing quantitative insights into thermodynamic properties such as binding affinities, solvation free energies, and conformational stability. Particularly, alchemical free energy simulation (AFES) methods have become important tools in structure-based drug design, where accurate estimation of relative or absolute binding free energies^1–4^ can guide hit discovery and lead optimization and thus reduce experimental overhead.

Multiple theoretical frameworks have been developed for performing free energy calculations. Thermodynamic integration (TI)^5^ remains one of the most widely adopted equilibrium-based methods, while non-equilibrium TI (NETI),^6^ grounded in Jarzynski’s equality, has gained popularity for its computational efficiency. Free energy perturbation (FEP),^7^ based on Zwanzig’s exponential averaging formula, has long been a foundational approach for calculating relative free energies, particularly in alchemical transformations. More recently, multi-site lambda dynamics (MSLD)^8^ has emerged as a promising approach for handling complex perturbation networks such as ligand series with multiple substitution sites.

In accordance with these theoretical frameworks, several software packages have implemented optimized engines for AFES. For example, Amber^9^ provides comprehensive support for the TI-based free energy calculation. GROMACS,^10^ in combination with external libraries such as *pmx*,^11,12^ enables the execution of NETI-based protocols. BLaDE,^8^ a GPU-accelerated platform, offers a dedicated implementation of MSLD, while Schrödinger’s *FEP+*^13^ and OpenMM-powered *OpenFE*^14^ provide widely used implementations of FEP for efficient relative free energy predictions.

While each method and simulation engine combination (Amber-TI, GROMACS-NETI, OpenMM-FEP, and BLaDE-MSLD) has been independently benchmarked,^8,9,14,15^ a systematic side-by-side comparison using the same molecular system and force field has not been yet reported due to unique implementations of these methods into different simulation engines, as well as the intended force field support. Addressing this challenge is further complicated by the lack of a unified tool to prepare the system and input files for all four methods in the same conditions. To overcome this limitation, we have developed *MolCube Alchemical Free Energy Simulator* (*MolCube-AFES*), an advanced module integrated into *MolCube-Apps* (https://apps.molcube.com), to streamline and standardize input generation for AFES across platforms. This study leverages *MolCube-AFES* to perform a comprehensive comparison of the four approaches on representative molecular systems, evaluating their accuracy, convergence behavior, computational efficiency, and practical usability. Our findings aim to provide valuable insights into the strengths, limitations, and optimal use cases of each method.

## 2. Methods

### 2.1 System Setup and Benchmark Targets

The protein-ligand complex structures used in this study were selected based on their relevance to the experimental ligand series. Specifically, we prioritized structures either directly derived from studies reporting binding affinities for the ligands of interest or those containing co-crystallized ligands with high structural similarity to the target compounds. Our study focused on the eight protein targets originally introduced by *Wang et al*.,^13^ the so-called JACS benchmark set widely used to assess AFES performance. To broaden the comparative scope, we additionally included a glucocorticoid receptor (GR) system described by *Weinstein et al.*^16^ and an HDAC6 inhibitor system from *Wang et al*.^17^ For the JACS benchmark targets, the initial protein-ligand complexes were obtained from experimentally determined structures deposited in the Protein Data Bank (PDB),^18^ with the following accession codes: BACE (4DJW),^19^ Tyk2 (4GIH),^20^ CDK2 (1H1Q),^21^ MCL1 (4HW3),^22^ JNK1 (2GMX),^23^ p38 (3FLY),^24^ thrombin (2ZFF),^24^ and PTP1B (2QBS).^25^ Because no co-crystal structure is available for the GR system, the protein-ligand complex was generated using Boltz-2,^26^ referencing PDB ID 3BQD for the binding-site conformation. For the HDAC6 inhibitor system, we used the docking model reported by *Wang et al.*,^17^ derived from the PDB ID 5EDU receptor structure, which served as our structural template in the absence of an experimentally determined complex.

To systematically and efficiently perform large-scale ΔΔG_bind_ calculations, we selected 10 transformations from each protein system in the JACS benchmark set (**Table S1**), ensuring that their ΔΔG_bind_ values span a relatively broad range based on available experimental data. Following the transformation-selection strategy described by *Han et al.*,^27^ this panel was constructed to capture the chemically diverse modifications typically explored during lead optimization in real-world drug discovery campaigns, including small substituent exchanges, ring additions or deletions, functional-group interconversions, and stereochemical changes. By incorporating both minor perturbations (e.g., hydrogen-to-halogen substitutions) and more substantial structural alterations, we aimed to assess the performance and generalizability of *MolCube-AFES* across a wide spectrum of chemically meaningful transformations.

The preparation of all simulation systems was streamlined through *MolCube-APPs* (**Figure 1**). Protein structures were prepared using the automated protocol in *PDB Reader*, which reconstructs any necessary missing residues, assigns protonation states assuming a pH of 7, and retains relevant crystallographic water molecules. Each system was solvated using *Solution Builder* in a cubic box with a 10 Å buffer of explicit water molecules and neutralized with 0.15 M KCl. The force field parameters were assigned using Amber ff19SB for proteins,^28^ OPC^29^ for water molecules in Amber-TI, GROMACS-NETI, and OpenMM-FEP and TIP3P^30^ in BLaDE-MSLD, and GAFF2^31^ with AM1-BCC charges^32^ for ligands. Note that BLaDE-MSLD does not support OPC water model, and the previous study by *Han et al.* shows minor influences of water models on ΔΔG_bind_.^27^

**Figure 1.**
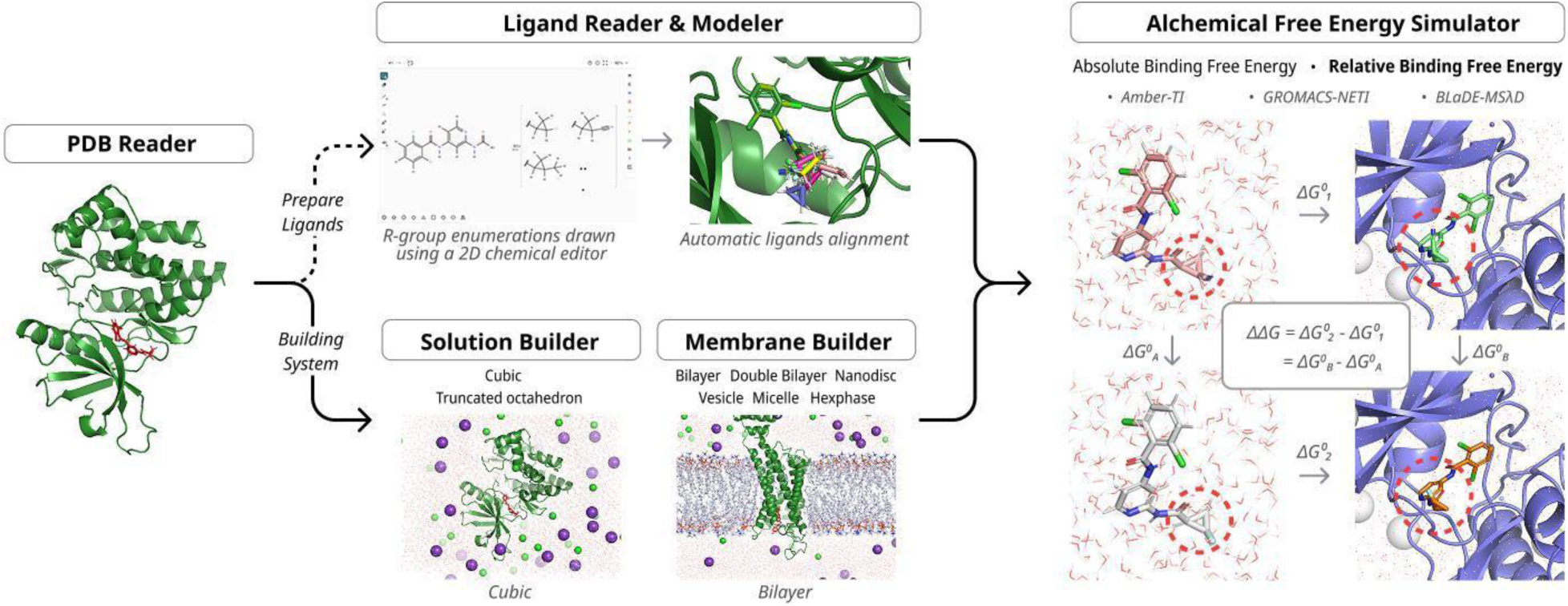
Schematic overview of the alchemical free energy simulation workflow implemented in *MOLCUBE-APPs*.

A series of congeneric ligands for each system were prepared using *Ligand Reader and Modeler (LRM)*, which (1) reads ligands from various sources, including uploaded SDF files or R-group enumerations drawn using a 2D chemical editor, (2) aligns them to the reference ligand from the corresponding *PDB Reader* project, and (3) assigns appropriate force field parameters. The procedure employs a maximum common substructure (MCS) algorithm^33^ to identify the shared atomic core, from which coordinates of the common atoms are directly transferred. A constrained conformation search is subsequently conducted for the remaining atoms, followed by a short energy minimization to yield a reasonable initial conformation. *LRM* automatically issues warnings for potential atomic overlaps, and users may manually adjust conformations if necessary. In this study, all systems were generated automatically, as potential overlaps are generally resolved during the minimization phase of the simulation.

*MolCube-AFES* then takes the prepared ligand sets and the solvated (or membrane) system to generate input files for Amber-TI, GROMACS-NETI, OpenMM-FEP, and BLaDE-MSLD relative binding free energy simulation, automating key steps in the preparation of perturbation networks and topologies. **Figures 2A-B** show screenshots of the *MolCube-AFES* user interface, illustrating a typical workflow for constructing TI calculation systems. Before generating the morph set, users may select ligands with a desired force field and choose the method to generate the perturbation network. For Amber-TI, GROMACS-NETI, and OpenMM-FEP, various types of perturbation networks are supported, including minimum spanning tree (MST), LOMAP,^34^ radial, and custom networks. Once a perturbation network is selected, the corresponding set of morph pairs can be generated automatically (**Figure 2A**). The resulting morphs can be viewed and modified in both tabular and graphical formats. In either case, users can find the corresponding core scaffold by hovering over the desired morph. A pairwise similarity matrix of ligands is also provided, allowing users to refine or update the generated network as needed. Once the set of morphs is determined, users can inspect and modify the core scaffold by clicking the core atoms in the 2D visualizer (**Figure 2B**, highlighted in red). In this study, we used custom networks for the JACS benchmark set and BMS-776532/BMS791826 analogues (GR, 3BQD) and TNI-97 series (HDAC6, 5EDU) for the other two case studies. The MCS and LOMAP network generation involves identifying the MCS for every ligand pair, scoring each MCS using the LOMAP algorithm,^34^ and assembling the perturbation network according to the selected strategy. Assigned morphs are realigned based on their pairwise MCS, which also defines the core scaffold of the hybrid topology later on. For Amber-TI, the MCS is further refined to exclude any heavy atom bonded to a hydrogen atom to account for SHAKE constraints. Hybrid single-dual topologies for Amber-TI and single topologies for GROMACS-NETI and OpenMM-FEP are then constructed, followed by solvation, neutralization, and removal of overlapping water molecules of both the ligand-only and protein-ligand complex systems.

**Figure 2.**
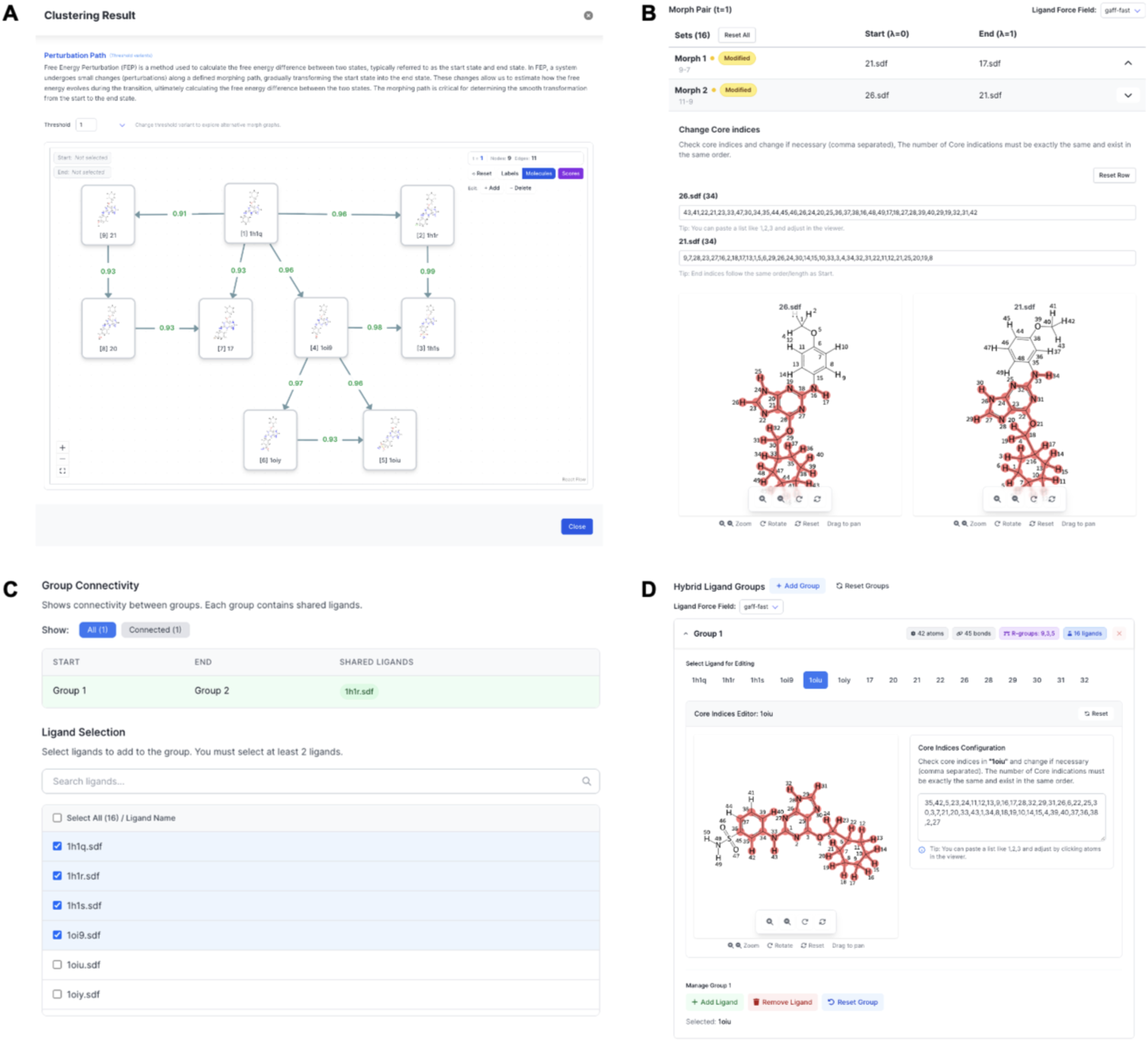
User interface of *MolCube-AFES*. TI-based calculation features of (A) automatically generated perturbation networks and (B) modification of core scaffold for each set of morphs. BLaDE-MSLD features of (C) selection of bridge ligand between groups, and (D) modification of group-wise MCS.

For BLaDE-MSLD, *MolCube-AFES* supports a ligand grouping option, which proceeds as follows: (1) the MCS of all ligands is computed to determine common cores across all ligands, (2) site-specific MCSs on substituents from (1) are calculated for every ligand pair, (3) ligands are grouped to maximize the common core with a user-defined maximum number of substituents per site, which can range from 2 to 9 (with a recommended value of 6). The resulting groups are *bridged* by sharing one common ligand to enable inter-group comparison of ΔΔG_bind_. In this study, we expanded the maximum number of substituents per site to 20 and evaluated the performance of various combinations of ligand groups, sites, and substituents. Additional factors affecting simulation efficiency, such as minimal recommended changes per group and substituent volume, among others, are also considered empirically. **Figures 2C-D** illustrate the *MolCube-AFES* user interface displaying both pre-grouped and grouped ligands for BLaDE-MSLD. The resulting ligand groups are then presented along with information, including the size of the core scaffold, the number of sites, and the number of substituents per site. Users may add, delete, or modify groups, ligands, or core atoms, as long as the groups maintain the *bridge* (**Figure 2C**). Single topologies are generated for each group by obtaining a group-wise MCS, realigning ligands accordingly, refining the MCS to remove atoms with differing bonded or non-bonded parameters, and performing charge renormalization (**Figure 2D**).^35^ The resulting topologies are then used to construct the ligand-only and protein-ligand complex systems, followed by solvation, neutralization, and removal of overlapping water molecules. Details of the preparation and simulation protocol for Amber-TI, GROMACS-NETI, OpenMM-FEP, and BLaDE-MSLD are provided in the subsequent sections.

### 2.2 Thermodynamic Integration with Amber

TI is a widely used equilibrium method for calculating free energy differences by integrating the ensemble average of the derivative of the system’s Hamiltonian (H) with respect to an alchemical thermodynamic coupling parameter λ. The parameter λ ranges from 0 (initial state) to 1 (final state), interpolating between two chemical species. The free energy change is computed via:

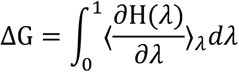

ΔΔG_bind_ between two ligands (L0 to L1) is calculated as:

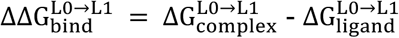

where 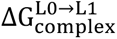 and 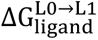 represent the free energy changes associated with the alchemical transformations of ligand L0 to L1 in the protein-ligand complex and in ligand-only solution, respectively.

To accurately perform TI calculations, particularly for systems undergoing significant chemical transformations, the underlying representation of the molecular states is critical. Amber employs a hybrid single-dual topology scheme in which both the initial and final ligand states are presented simultaneously within the system with shared coordinates for common scaffold atoms, and their interactions with the environment are modulated by λ. This approach incorporates the advantages of a dual topology framework, which maintains separate, non-interacting topologies for each state to ensure structural and energetic consistency throughout the transformation. As a result, it is particularly effective in handling complex chemical changes such as mutations or ligand modifications, while enabling accurate estimation of relative free energies by minimizing artifacts from topological discontinuities.

All Amber-TI molecular dynamics (MD) simulations were performed under periodic boundary conditions (PBCs) in the isothermal-isobaric (NPT) ensemble at 300 K and 1 atm. Temperature was maintained using Langevin dynamics with a collision frequency of 5 ps⁻¹, while pressure was regulated with a Monte Carlo barostat.^36^ To enable the use of a 4-fs integration time step, covalent bonds involving hydrogen atoms were constrained using SHAKE algorithm, and hydrogen mass repartitioning (HMR) was applied.^37^ Nonbonded interactions were truncated at 9 Å. For simulations involving alchemical transformations, atoms undergoing modification were treated using a softcore potential,^9^ and van der Waals and electrostatic interactions in these regions were smoothly scaled during the transformation.

Prior to dynamics, all systems were subjected to energy minimization to remove unfavorable contacts. Minimization was carried out using a steepest descent algorithm for up to 1,000 steps. The softcore potential was applied during this stage to the transforming regions of the alchemical perturbation, and bond constraints were selectively removed from these atoms. Following minimization, the systems were gradually heated from 1 K to 300 K over a 5-ps period using a linear temperature ramp. Heating was performed under NPT conditions using Langevin dynamics and the Monte Carlo barostat. This step ensured proper equilibration of the system prior to production. During heating, atoms involved in the alchemical transformation were excluded from bond constraints to allow for flexible structural adjustment.

Production simulations for Amber-TI were carried out using the hybrid single-dual topology approach. Atoms present only in the initial or final ligand state were explicitly defined, and the softcore potential was applied throughout the transformation to avoid singularities. Each alchemical transformation was simulated across 12 discrete λ values (0.0000, 0.0479, 0.1151, 0.2063, 0.3161, 0.4374, 0.5626, 0.6839, 0.7937, 0.8850, 0.9521, and 1.0000).^38^ For each λ window, a 5-ns MD simulation was performed. To assess statistical variability, four independent replicates were conducted per transformation, and the average ΔΔG_bind_ value was reported.^39^ During each simulation, energy evaluations for all λ states were recorded to facilitate multistate free energy estimation using the multistate Bennett acceptance ratio (MBAR) method.^40^

### 2.3 Non-equilibrium Thermodynamic Integration with GROMACS

The GROMACS-NETI method leverages non-equilibrium dynamics and Jarzynski’s equality to estimate free energy differences from repeated fast switching simulations between initial and final states. The core idea is to generate multiple forward and reverse transformations, from which the free energy difference is extracted via exponential averaging:

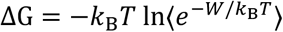

where *k*_B_is the Boltzmann constant, *T* is temperature, *W* is the work done on the system during the non-equilibrium transformation, and the average is taken over many trajectories. This method avoids the need to sample intermediate λ-states, potentially reducing a simulation cost.

To enable these transformations between states, GROMACS employs a single topology approach, which provides a consistent molecular representation throughout the simulation. In this framework, both the initial (L0) and final (L1) molecules are encoded within a single topology: common scaffold atoms to both states are shared, while perturbed atoms are treated as "dummy" atoms that are gradually turned on or off using a coupling parameter λ. This λ parameter modulates the interactions (e.g., charges, van der Waals, bonded terms) of the atoms across a series of virtual states, allowing a smooth transformation between L0 and L1. The single topology method ensures a continuous transformation pathway and avoids complications from abrupt changes in atomic composition.

All NETI simulations were carried out using GROMACS 2025.2 with the PBCs applied in all three dimensions. All simulations were performed in the NPT ensemble at 300 K and 1 bar. Temperature was regulated using the velocity-rescaling thermostat with a stochastic term (v-rescale), and pressure was maintained using the stochastic cell rescaling barostat (c-rescale). A 4-fs time step together with HMR was used, with all covalent bonds involving hydrogen atoms constrained to enable stable integration. Prior to alchemical simulations, all systems were subjected to energy minimization using the steepest descent algorithm to remove steric clashes and relax high-energy bad contacts. Following minimization, a two-stage equilibration protocol was applied. The system was first equilibrated under constant volume and temperature (NVT) conditions and then under NPT, with each phase conducted for 500 ps and 10 ns, respectively. During NVT equilibration, positional restraints were applied to heavy atoms to stabilize the system and preserve the overall structure.

Non-equilibrium alchemical transformations were performed with simulations initiated from equilibrated configurations at the two end states (λ = 0 and λ = 1). Alchemical regions were defined explicitly, and softcore potentials were applied to avoid singularities during the transition. Each transformation was executed using a linear pull of the λ-parameter over 250 ps, with 100 replicates forward and reverse runs to ensure statistical convergence. Four independent trials were performed for each direction to improve statistical reliability. Free energy differences were estimated from the work distributions obtained in forward and reverse transformations. Post-processing was conducted using the *pmx* v.4.0.3 toolkit,^11,12^ which generated and analyzed the nonequilibrium work trajectories to compute the final free energy estimates.

### 2.4 Replica Exchange Free Energy Perturbation with OpenMM

Similar to TI, alchemical FEP is a statistical mechanical approach for computing free energy differences between two thermodynamic states by introducing a nonphysical alchemical pathway that continuously connects them. Using this framework, relative binding free energies between two ligands, L0 and L1, were computed by transforming ligand L0 into ligand L1 in the bound state. The relative free energy difference is defined as

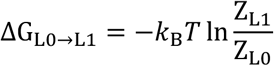

where *k*_B_ is the Boltzmann constant, T is the temperature, and Z_L0_ and Z_L1_ are the configurational partition functions corresponding to the fully interacting states of ligands L0 and L1, respectively. Direct evaluation of this expression is impractical due to limited phase-space overlap between the end states; therefore, an alchemical coupling parameter λ is introduced to define a continuous transformation between the two states. Along this alchemical pathway, the system Hamiltonian is defined as

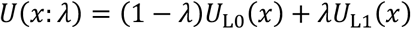

where *U*_L0_(x) and *U*_L1_(x) denote the potential energy functions of the initial and final ligand states, respectively, and x represents the full set of atomic coordinates. The free energy difference is obtained by sampling a discrete set of intermediate alchemical states λ_i_ spanning the interval from λ = 0 to λ = 1.

Efficient sampling of intermediate λ-states is essential for accurate free energy estimation but is often hindered by slow conformational relaxation and poor configurational overlap between neighboring states, particularly near the end points of the alchemical transformation. To address these challenges, Hamiltonian replica exchange can be employed. In this approach, multiple replicas corresponding to different λ-states are simulated in parallel, and periodic exchange attempts of configurations between neighboring replicas are performed. Exchanges are accepted according to the Metropolis criterion,

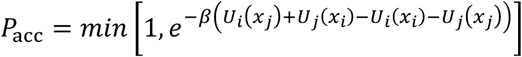

where *β = 1/k*_B_T, *U*_i_ and *U*_j_ denote the Hamiltonians associated with λ_i_ and λ_j_, and x_i_ and x_j_ are the configurations of replicas i and j, respectively. This exchange scheme enables configurations to diffuse efficiently across alchemical space, thereby improving overlap between adjacent λ-states and accelerating convergence.

All alchemical FEP calculations were performed using OpenMM within a single-topology framework. For each ligand transformation, a hybrid system was constructed in which the initial and final ligand states were combined into a single alchemical representation. Interactions corresponding to the two ligand states were smoothly interpolated as a function of λ, ensuring a continuous transformation while maintaining a physically consistent description of the system throughout the simulation. Hybrid systems were generated using an in-house Python workflow inspired by the *OpenFE* framework, without directly relying on the *OpenFE* library.

Hamiltonian replica exchange simulations were carried out using the alchemical utilities provided by *openmmtools.alchemy*.^41^ A series of intermediate λ-states was defined between the fully interacting initial and final ligand states. At each λ, an independent replica was propagated using Langevin dynamics at 300 K with a timestep of 2 fs. Exchange attempts between neighboring λ-replicas were attempted every 2.5 ps. Each replica was equilibrated for 1 ns prior to production sampling. Following equilibration, production simulations of 5 ns per λ-state were performed. Configurations collected from all replicas were used for free energy analysis.

Relative free energy differences were estimated using the MBAR method. MBAR analysis was performed using the implementation available in *openmmtools.alchemy*, yielding statistically optimal free energy estimates by combining information from all λ-states. Convergence was assessed by examining overlap of reduced potential energy distributions between adjacent λ-states.

### 2.5 Multi-Site Lambda Dynamics with BLaDE

BLaDE supports MSLD, a dynamic alchemical approach in which λ-parameters are treated as additional degrees of freedom that evolve alongside atomic positions during simulation. Unlike traditional free energy methods such as FEP or TI, which have fixed λ values and require separate simulations for each state, MSLD enables the system to dynamically explore multiple alchemical states within a single trajectory. This makes BLaDE particularly effective for evaluating large combinatorial libraries of ligands with multiple substitution sites.

BLaDE adopts a single topology scheme. Unlike GROMACS, however, it does not support assigning multiple parameter sets, such as atom type, mass, and charge, to a single atom. This limitation introduces the need for additional steps, including core scaffold refinement and charge renormalization, to ensure accurate representation of atomic interactions. An initial core scaffold could be identified based on atomic connectivity and element type. However, the mapped core atoms may not always share identical force field parameters. To address this issue, in this study, the core scaffold is refined by excluding atoms with different bonded or non-bonded parameters, ensuring consistency within the core. Following the core scaffold refinement, charge renormalization is performed to maintain consistent core atom charges across ligands. For each core atom, partial charges are first extracted from all ligands and averaged. These averaged values are then reassigned to the corresponding core atoms. To preserve the original net charge of each molecule, the charges of the site (non-core) atoms are adjusted accordingly, ensuring overall charge neutrality and force field compatibility.^35^

To ensure accurate sampling and efficient convergence, BLaDE employs a biasing potential^42^ and adaptive landscape flattening (ALF) scheme^43,44^ that promote thorough exploration of alchemical space. A biasing potential flattens the free energy landscape and reduces barriers between alchemical states, and an adaptive scheme continuously updates this bias to promote transitions into under-sampled regions of λ-space, preventing the system from becoming trapped in low-energy basins and enabling uniform exploration across the combinatorial ligand set. Together, these techniques enhance the efficiency of sampling across a high-dimensional combinatorial space of substituents and ensure that free energy differences can be estimated with minimal statistical uncertainty.

Free energy differences are computed based on the time-averaged populations of alchemical states sampled throughout the trajectory, using reweighting techniques to correct for bias and ensure statistically robust estimates. The relative free energy between two alchemical states L0 and L1 is calculated using:

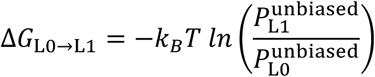

where 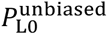 and 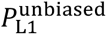 are the reweighted, time-averaged populations of states L0 and L1, respectively. These were obtained by correcting the raw (biased) state populations 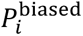 using the biasing potential *V*_bias_(*λ_i_*) applied during the simulation:

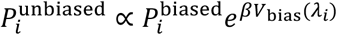

This reweighting ensures that the free energy estimates reflect the true underlying thermodynamic distribution, independent of the enhanced sampling strategy used to accelerate convergence. Therefore, the resulting free energies are statistically robust and account for uneven sampling due to adaptive biasing.

All BLaDE-MSLD simulations were conducted using *ALF* package^43^ with BLaDE^8^ as the MD engine, except during the energy minimization and short equilibration phase. Energy minimization and short equilibration was conducted using OpenMM (version 8.2.0) under the PBCs in the NPT ensemble at 300 K and 1 bar. Temperature was regulated using Langevin dynamics with a friction coefficient of 1.0 ps⁻¹ and a time step of 2 fs, while pressure was maintained via a Monte Carlo barostat. Long-range electrostatics interactions were treated using the Particle Mesh Ewald (PME) method^45^ with a 9 Å non-bonded cutoff. All bonds involving hydrogen atoms were constrained to their equilibrium lengths. The system was minimized for up to 5,000 steps, followed by a 500-ps equilibration run employing simulated annealing to ensure adequate structural relaxation prior to subsequent ALF-based sampling.

Following energy minimization and short equilibration using OpenMM, all systems underwent a hierarchical ALF-based simulation protocol. Steps 1-111 comprised the flattening phase, consisting of 100 cycles of 26-ps equilibrium and 78-ps production followed by 11 cycles of 0.25-ns equilibrium and 0.75-ns production to iteratively refine the bias parameters. Step 112 consisted of 5-ns equilibration simulations with five independent trials (steps 112a-e), while Step 113 represented the production phase, consisting of 40-ns simulations for the protein-ligand complex (two trials; steps 113a,b) and 20-ns simulations for the ligand alone (five trials; steps 113a-e). Each production run was followed by post-processing analysis to estimate the free energy change upon chemical perturbation using a histogram-based estimator with the λ cutoff parameter set to 0.

The BLaDE-MSLD simulation set up for λ-dynamics followed the general protocol of *ALF*.^43^ All simulations were performed under the PBCs within a cubic box in the NPT ensemble at 300 K and 1 atm. Temperature was maintained using a Langevin thermostat with a friction coefficient of 0.1 ps⁻¹, while pressure was controlled using a Monte Carlo barostat^36^ with a maximum volume fluctuation of 100 Å³. The Leapfrog Verlet integrator was used with a 2-fs integration time step. Electrostatic interactions were computed using PME^45^ with an interpolation order of 6, Ewald coefficient of 3.125 Å⁻¹, and a grid spacing of 1.1 Å. Non-bonded interactions were truncated at 9 Å. Hydrogen-containing bond constraints were enforced using the SHAKE algorithm. A softcore potential was applied during the initial 14 steps and deactivated thereafter. The ALF biasing function employed an exponential constant (λ_nexp_) of 5.5. Production runs were conducted under the same conditions as bias optimization. Note that, unlike Amber-TI and GROMACS-NETI, BLaDE-MSLD simulations in this study, we could not use HMR or the OPC water model due to the limitation of single topology scheme and the challenges associated with incorporating virtual sites in the current BLaDE implementation.

All BLaDE-MSLD simulations were executed within a single *ALF* project per system, with the exceptions of p38 and PTP1B. Since *ALF* natively supports up to 10 substituents per site, we recomputed the implicit-constraint free energy term G_imp_ to accommodate up to 20 substituents, thereby allowing every benchmark system to be represented as a single multi-ligand topology project. Both p38 and PTP1B systems with the highest MUE were simulated using two (PTP1B) and three (p38) separate *ALF* projects to assess potential performance improvements.

### 2.6 Evaluation Metrics

To quantify agreement between predicted and reference ΔΔG_bind_, we report both absolute error and correlation-based measures, computed as per-target (per-protein) statistics averaged over the eight protein systems in the JACS benchmark. Root-mean-square error (RMSE) and mean unsigned error (MUE; equivalent to mean absolute error, MAE) measure absolute deviation between predicted and reference ΔΔG_bind_ values in kcal/mol. Lower RMSE/MUE indicates better absolute agreement with the reference. The coefficient of determination (R²) and Pearson’s correlation coefficient (r_p_) evaluate whether a method preserves the relative ordering of ΔΔG_bind_ values across transformations. Two correlation coefficients reflect target-specific structure-activity relationship trends. Finally, Acc ±1 and Acc ±2 report the fraction of predictions whose absolute error relative to the reference falls within ±1 kcal/mol or ±2 kcal/mol, respectively. These accuracy-in-window metrics provide a practical assessment of model performance by quantifying the proportion of affinity changes predicted within ranges that is generally regarded as informative for decision-making in compound design. Here again, values are averaged on a per-protein basis to prevent any single target from disproportionately influencing the results. Together, these metrics allow us to distinguish between two desirable aspects of performance: low absolute error (RMSE, MUE, Acc ±1, and Acc ±2), which demonstrates how close the predicted values are to the experimental measurements in terms of their magnitude, and high correlation (R², r_p_), which indicates how consistently the method preserves the correct binding ranking of ligand modifications within a series.

### 2.7 Bootstrap Statistical Analysis

To assess the statistical uncertainty of the performance metrics and the significance of pairwise method differences, we applied two complementary bootstrap resampling procedures (2,000 iterations each). For each method, 95% bootstrap confidence intervals for MUE were estimated by simple non-parametric bootstrap resampling. In each iteration, 80 morphs (JACS benchmark set) were drawn with replacement from the full dataset, and MUE was computed on the resample. The 2.5^th^ and 97.5^th^ percentiles of the resulting distribution were taken as the 95% confidence interval. To test whether any two methods differ significantly in MUE, we used a paired bootstrap procedure. In each iteration, the same set of 80 morphs, drawn with replacement, was used to compute MUE for both methods simultaneously, and the difference in MUE (ΔMUE = MUE_A_ – MUE_B_) was recorded. Resampling morphs in pairs preserves the morph-level correlation between the methods, which ensures that the difficulty of individual transformation affects the comparison. A one-sided p-value was computed as the fraction of bootstrap samples in which the observed direction of the difference was not reproduced (e.g., fraction of resamples where ΔMUE ≥ 0).

### 2.8 Ensemble Combination Strategies

To evaluate whether combining predictions from multiple methods improves accuracy, we tested four ensemble strategies applied to the mean predictions (averaged across four replicas) from each method. Equal weight averaging computes the ensemble prediction as the unweighted mean of the four method predictions. Inverse-MUE weighting assigns each method a weight inversely proportional to its global MUE (*w_i_* = (∑*_j_* MUE*_j_*)⁄MUE*_i_*), so that more accurate methods contribute more to the ensemble prediction. Weights were computed from the global MUE of each method over all 80 morphs. Oracle ensemble represents an idealized upper bound on per-protein selection; for each protein, the method with the lowest MUE on that protein is selected, and its predictions are used exclusively. This is not a deployable strategy (it requires knowledge of the ground truth), but serves as a reference for the maximum benefit achievable by perfect protein-level method selection. Linear stacking with leave-one-protein-out cross-validation (LOPO-CV) fits a linear combination of the four method predictions using ordinary least squares, with weights constrained to be non-negative and sum to one. The weights were estimated using LOPO-CV to avoid overfitting on the 80-morph dataset. In each fold, all morphs from one protein were held out, weights were fit on the remaining seven proteins, and predictions for the held-out protein were generated using those weights. This procedure was repeated for each of the eight proteins, and the resulting held-out predictions were used to compute the ensemble MUE. LOPO-CV ensures that the weights used to predict each protein were never trained on data from that protein, providing a realistic estimate of generalization performance.

## 3. Results and Discussion

### 3.1 JACS Benchmark

#### 3.1.1 Accuracy of Ligand Binding Free Energy Predictions

The performance of four alchemical free energy approaches (Amber-TI, GROMACS-NETI, OpenMM-FEP, and BLaDE-MSLD) was evaluated on the JACS benchmark consisting of eight protein targets and ten ligand transformations per target (80 total transformations in **Table S1**). For each method, the predicted ΔΔG_bind_ were compared against the corresponding experimental binding affinity. Performance was assessed using two complementary views, namely scatter plots, which visualize how closely the predicted values track the reference values (**Figure 3**), and tabulated statistical metrics, which summarize per-target agreement (**Table 1**). Note that the prediction results are the average ones from 4 independent runs in this study (see the raw data in **AFES_Supplementary2.xlsx**).

**Figure 3.**
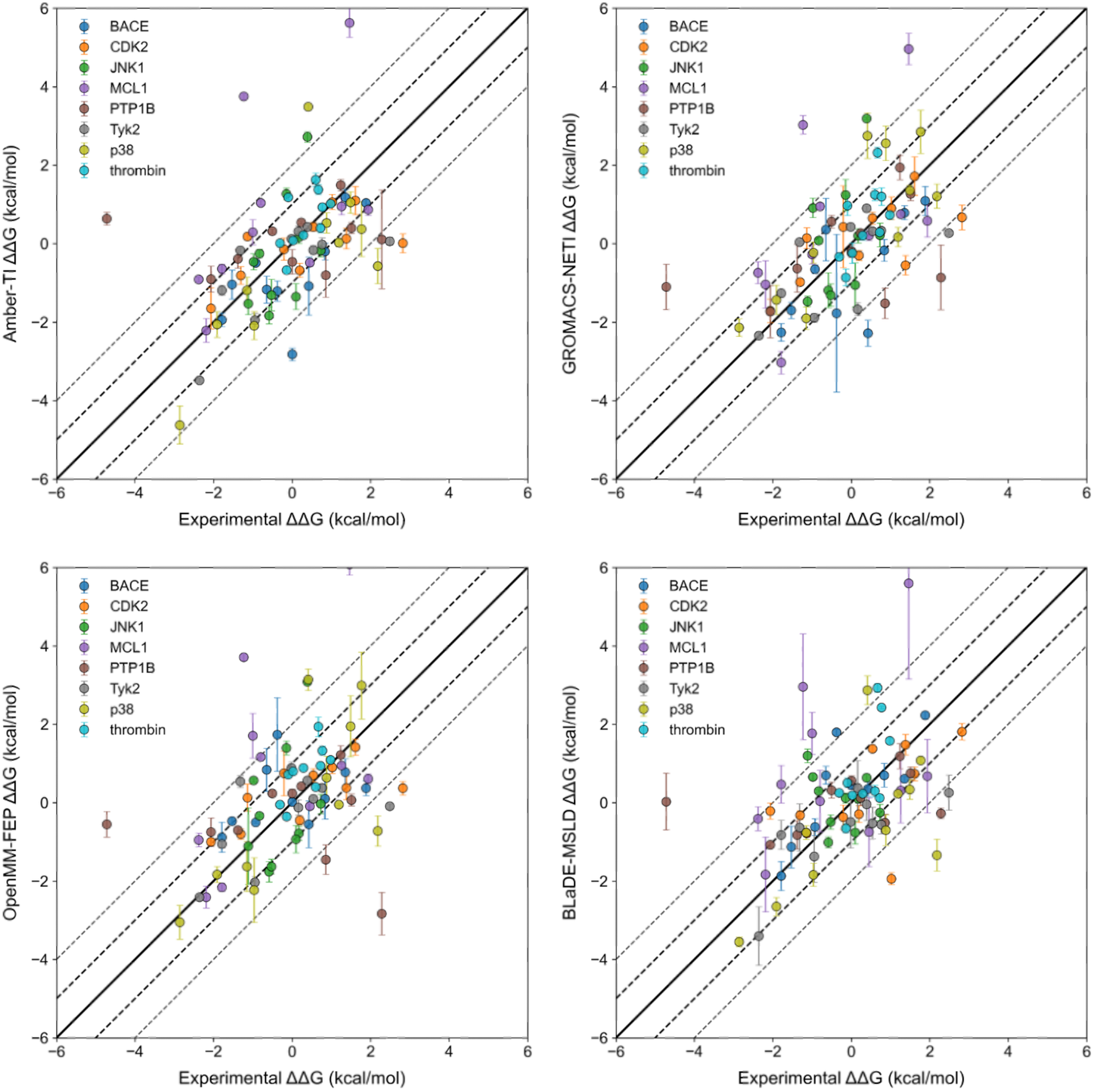
Comparison of ΔΔG_bind_ prediction using *MolCube-AFES* (Amber-TI, GROMACS-NETI, OpenMM-FEP, BLaDE-MSLD) with the experiment results.

**Table 1.** Comparison of ΔΔG_bind_ values across the four methods with the experimental results.

| | RMSE<br>(kcal/mol) | MUE<br>(kcal/mol) | Correlation $R^2$ | $r_P$ | Acc $\pm 1$ | Acc $\pm 2$ |
| --- | --- | --- | --- | --- | --- | --- |
| **Amber-TI** | 1.37 | 1.03 | 0.35 | 0.56 | 0.59 | 0.88 |
| **GROMACS-NETI** | 1.28 | 0.99 | 0.38 | 0.61 | 0.61 | 0.88 |
| **OpenMM-FEP** | 1.44 | 1.11 | 0.30 | 0.48 | 0.59 | 0.85 |
| **BLaDE-MSLD** | 1.38 | 1.07 | 0.33 | 0.47 | 0.64 | 0.84 |

The ΔΔG_bind_ comparison scatter plots show that all four predictions cluster near the x=y line with 59-64% of transformations within ±1 kcal/mol and 84-88% within ±2 kcal/mol from experiment, indicating that all approaches capture both the direction and approximate magnitude of the binding affinity changes (**Figure 3**). Among the four methods, GROMACS-NETI shows the lowest RMSE and MUE (1.28 and 0.99 kcal/mol; Acc ±2 = 0.88), followed by Amber-TI (1.37/1.03 kcal/mol; Acc ±2 = 0.88), BLaDE-MSLD (1.38/1.07 kcal/mol; Acc ±2 = 0.84), and OpenMM-FEP (1.44/1.11 kcal/mol; Acc ±2 = 0.85).

All four methods produce extreme outliers particularly for MCL1 and PTP1B, as evident from the scatter plots, implying a higher risk of occasional large errors; also see **Table S2**. In the case of MCL1, the apparent outliers can be attributed to alchemical transformations that involve large structural perturbations, such as O → N atom substitutions within a fused 5.6-bicyclic ring or transformations of a benzene ring into a biaryl system (Amber-TI RMSE = 2.30 kcal/mol, MUE = 1.72 kcal/mol; GROMACS-NETI RMSE = 2.04 kcal/mol, MUE = 1.64 kcal/mol; OpenMM-FEP RMSE = 2.48 kcal/mol, MUE = 1.84 kcal/mol; BLaDE-MSLD RMSE = 2.37 kcal/mol, MUE = 1.99 kcal/mol). Similarly, for PTP1B, the observed outliers appear to arise from morphs that involve large structural perturbations, for example, when pronounced substituent rearrangements occur at a ring nitrogen, particularly in transformations introducing sulfur-containing, N-linked substituents, or when an additional seven-membered ring is introduced. Outliers are also associated with structurally small but electronically significant transformations, e.g., ring-bound H → Cl substitutions (Amber-TI RMSE = 2.02 kcal/mol, MUE = 1.43 kcal/mol; GROMACS-NETI RMSE = 1.76 kcal/mol, MUE = 1.25 kcal/mol; OpenMM-FEP RMSE = 2.32 kcal/mol, MUE = 1.62 kcal/mol; BLaDE-MSLD RMSE = 1.84 kcal/mol, MUE = 1.26 kcal/mol).

The numbers of ligands, sites, and substitutions used in the BLaDE-MSLD simulations are summarized in **Table S3**. Among the BLaDE-MSLD results, PTP1B exhibited particularly poor accuracy in its single-project form (**Table S4** 1 group MUE = 12.17 kcal/mol). Inspection indicated that the perturbations among its substituents were excessively heterogeneous, ranging from minimal fragments such as a methyl group to bulky bicyclic groups bridged by a sulfonamide, resulting in transformations that were not physically traversable within a single alchemical simulation (**Figure 4**). To address this, we performed multi-group simulations on the two worst-performing systems, p38 and PTP1B (**Figure 4** and **Table S3**). **Table S4** reports the computed ΔΔG_bind_ values for each morph in the p38 and PTP1B systems. For PTP1B, dividing the ligands into two groups according to their volumetric and topological disparity led to a dramatic improvement (MUE 12.17 → 1.26; r_p_ −0.11 → 0.36 in **Tables S2, S4**). For p38, partitioning the ligands into three groups based on substituent site resulted in a modest gain (MUE 1.54 → 1.30; r_p_ 0.23 → 0.65 in **Tables S2, S4**).

**Figure 4.**
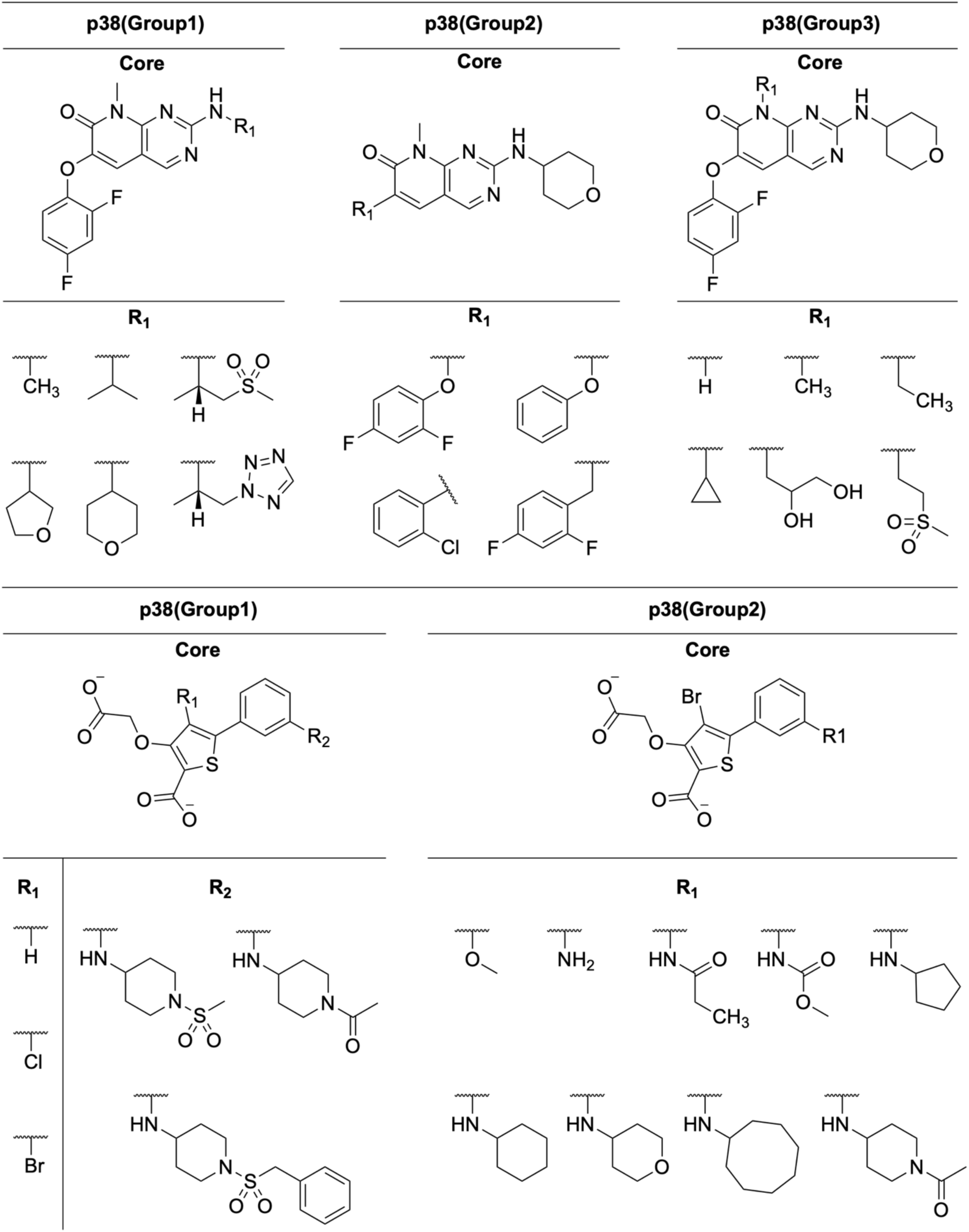
p38 (group 1, 2, and 3) and PTP1B (group 1 and 2) ligand structures with sites and substitution groups.

Among the four methods, GROMACS-NETI achieves the best quantitative agreement with experiment across all metrics, attaining the lowest RMSE (1.28 kcal/mol) and MUE (0.99 kcal/mol) alongside the highest Acc ±1 (0.61) and Acc ±2 (0.88, tied with Amber-TI), and shows a relatively uniform distribution of errors with few extreme outliers. BLaDE-MSLD, using the multi-group simulation strategy for p38 and PTP1B, achieves RMSE = 1.38 kcal/mol and MUE = 1.07 kcal/mol with Acc ±1 = 0.47 and Acc ±2 = 0.64, broadly comparable to OpenMM-FEP and Amber-TI.

All four methods exhibit statistically meaningful correlations with experiment, indicating that they capture the underlying structure-activity trends across the dataset. BLaDE-MSLD and OpenMM-FEP show the lowest correlations (R² = 0.33, rₚ = 0.47 and R² = 0.30, rₚ = 0.48, respectively), with BLaDE-MSLD showing compressed dynamic range particularly for targets such as p38 and JNK1, and OpenMM-FEP exhibiting sensitivity to a few larger outliers. Amber-TI achieves moderate correlation (R² = 0.35, rₚ = 0.56). GROMACS-NETI attains the highest correlation coefficients (R² = 0.38, rₚ = 0.61), reflecting more consistent preservation of rank ordering across ligand modifications. Taken together, GROMACS-NETI provides the best overall combination of quantitative accuracy (lowest RMSE/MUE) and rank-ordering performance (highest R², rₚ, and Kendall’s τ = 0.44) across the JACS benchmark. The performance differences among the methods, however, are modest and their statistical significance is evaluated more rigorously in Section 3.1.4.

In addition to our internal evaluation, we also examined the agreement with the results reported by *Wang et al.* that used a widely adopted commercial workflow, *Schrödinger FEP+* (**Figure S1** and **Table S5**). The RMSE and MUE reported by *Wang et al.* (1.23/0.99 kcal/mol) are comparable to those of GROMACS-NETI (1.34/0.99 kcal/mol). Their Acc ±1 value (0.57) is similar to those from our four methods (0.59-0.64), whereas Acc ±2 (0.89) is nearly identical to that of GROMACS-NETI and Amber-TI (both 0.88). In terms of correlation, *Wang et al.* achieved R² = 0.40 and rₚ = 0.62, somewhat higher than those of all four methods in this study (R² = 0.30-0.38). Overall, the commercial *FEP+* workflow and the open-source methods evaluated here achieve broadly comparable accuracy on the JACS benchmark, with GROMACS-NETI offering the closest performance to *FEP+* among our implementations.

#### 3.1.2 Convergence Analysis

For the Amber-TI protocol, we performed convergence analysis to ensure that results obtained with 12 λ windows and 5 ns per window reflect equilibrated sampling. The λ-window overlap matrix and the forward and reverse ΔΔG_bind_ for the 6e → 6b transformation of thrombin are shown in **Figures 5A**. As reported by *Klimovich et al.*,^46^ agreement between forward and reverse estimates within their uncertainties indicates a convergence of ΔΔG_bind_; in our case, the two estimates are statistically consistent under the 12-window protocol. Previous studies further suggest that the λ-window overlap matrix should be at least tridiagonal, with diagonal and near-diagonal elements with ≥ 0.03, in order to obtain trustworthy free energies.^46^ This expected minimum-overlap structure is observed in our calculations, supporting both numerical stability and physical reliability of the TI-based calculations.

**Figure 5.**
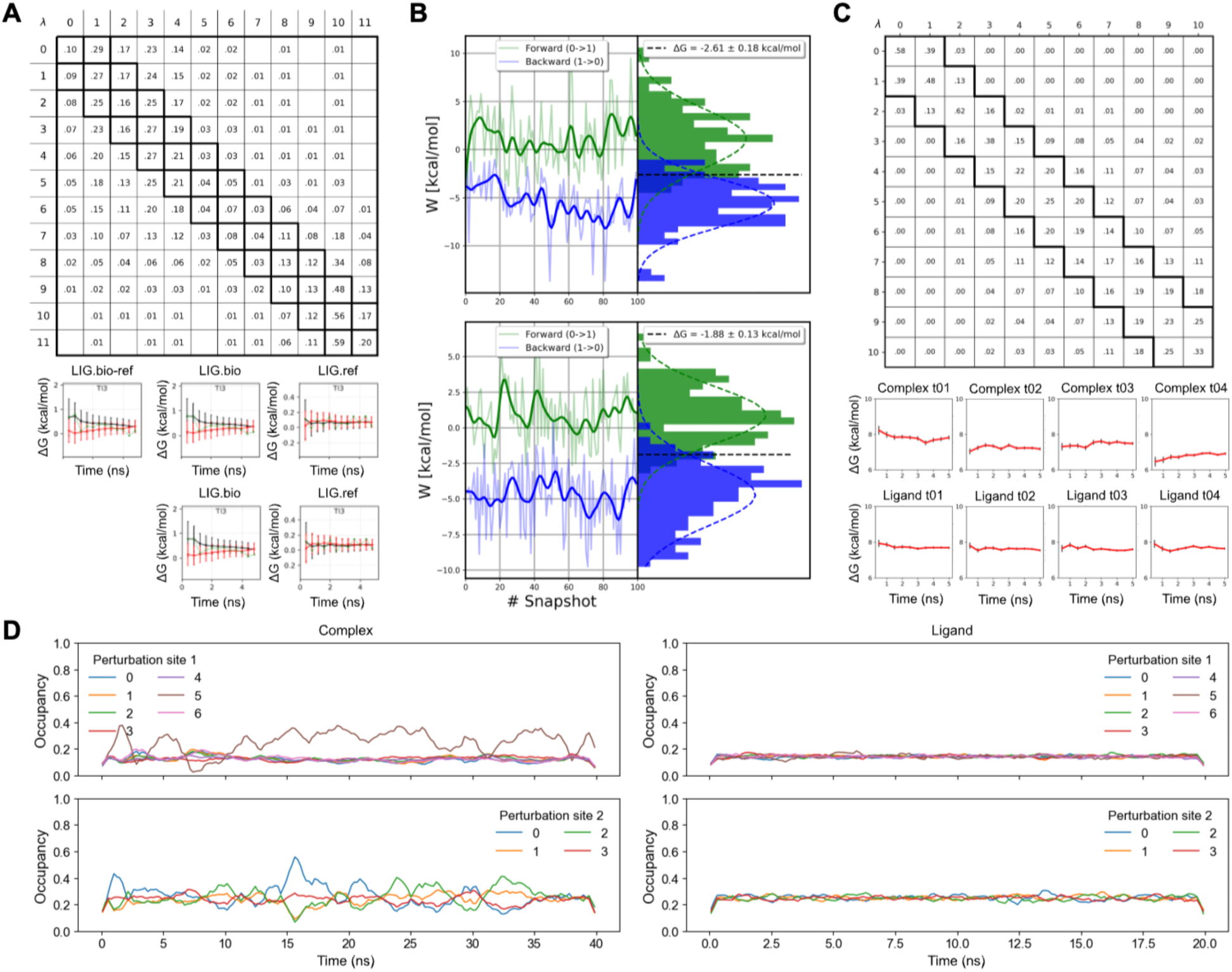
Sampling quality, convergence behavior, and state-space exploration evaluated using Amber-TI, GROMACS-NETI, OpenMM-FEP, and BLaDE-MSLD. (A) Amber-TI analysis results using a 12-window, 5 ns per window protocol: the λ-window overlap matrix and the time evolution of the forward (black solid line), reverse (red solid line), and block-averaged (green dotted line) estimates of ΔG_bind_ for each λ value. (B) GROMACS-NETI *pmx* analysis results: complex and ligand states corresponding forward (green) and reverse (blue) work distributions (bold lines for the mean work as a function of snapshot) together with the estimated free energy change and statistical uncertainty. (C) OpenMM-FEP replica exchange analysis using a 12-window, 5 ns per window protocol: overlap matrix using MBAR and the time evolution of ΔG_bind_ estimates for each λ value. (D) BLaDE-MSLD transition analysis: λ profiles at site 1 and site 2 of the thrombin system (complex and ligand) reflect ligand occupancy during the simulation.

In GROMACS-NETI, ΔG_bind_ between two alchemical states L0 and L1 is estimated separately for the complex and for the isolated ligand using nonequilibrium switching trajectories. For each transformation, the external work applied to drive the system from L0 → L1 (forward) or L1 → L0 (reverse) is integrated, yielding forward and reverse work ensembles, W_f_ and W_r_, from which *pmx* derives ΔG and its associated uncertainty. In **Figures 5B**, the forward (green) and backward (blue) nonequilibrium switching trajectories are shown together with bold lines indicating the mean work as a function of snapshot. The corresponding forward and backward work histogram are also presented, and the statistics of their overlap region is used to estimate ΔG, so that extent of this overlap provides a practical indicator of convergence. The two work distributions exhibit a substantial overlap and yield a well-defined crossing point, consistent with the Crooks fluctuation theorem,^47^ producing ΔG estimates with reported uncertainties on the order of ∼0.13-0.18 kcal/mol in these examples. The absence of heavy tails or strong skew further suggests that the switching is not excessively dissipative and that the trajectories are well behaved. Taken together, these diagnostics indicate that the NETI calculations for both the complex and the ligand are well converged and numerically robust under our simulation protocol.

For OpenMM-FEP, we analyzed the replica-exchange simulations using *openmmtools*. To assess whether neighboring λ windows sample sufficiently similar configurations, we computed an MBAR-based per-λ overlap matrix (**Figure 5C**). Briefly, we collected configurations generated during replica exchange and extracted the corresponding reduced energies evaluated across all λ states. These multi-state energy data were then combined using MBAR to quantify the extent to which configurations sampled at one λ can be reweighted to other λ states. The resulting overlap matrix provides a simple diagnostic of sampling continuity along λ, with large diagonal elements as expected, appreciable values between adjacent λ states indicating good window-to-window connectivity, and progressively smaller values for more distant λ pairs reflecting increasingly distinct ensembles. Overall, the pattern in **Figure 5C** supports adequate overlap between neighboring windows and suggests that the OpenMM-FEP free-energy estimates are not driven by gaps in sampling along λ.

BLaDE-MSLD employs a multisite λ-dynamics framework that enables simultaneous sampling of multiple substituents across predefined modification sites within a single simulation. Instead of performing pairwise alchemical transformations between ligand variants, substituent states are continuously modulated through λ-like order parameters that dynamically evolve along the trajectory. To characterize the sampling behavior, we monitored the time evolution of the effective occupancy of each substituent at individual sites (**Figure 5D**). Each trace represents the population (i.e., occupancy) of a given substituent state during the simulation. The distribution of occupancies provides direct insight into the thermodynamic and kinetic behavior of the system. When multiple substituents exhibit comparable occupancies, this indicates either small free-energy differences between states or continuous sampling across substituent space, reflecting efficient exploration of the available chemical configurations. In contrast, when a single substituent dominates the occupancy, this suggests a clear thermodynamic preference corresponding to a lower free-energy state. At the same time, the temporal fluctuations of the occupancies reveal the extent of dynamic sampling. Frequent changes in occupancy indicate that the system transitions between substituent states on the simulation timescale, whereas a lack of fluctuation would suggest kinetic trapping in a particular configuration. Therefore, analysis of substituent occupancies allows simultaneous assessment of thermodynamic weighting (through relative populations) and kinetic accessibility (through temporal fluctuations), providing a comprehensive picture of sampling behavior in BLaDE-MSLD simulations.

#### 3.1.3 Statistical Robustness of Performance Differences

With the well-converged ΔG_bind_ from the four AFES methods, to assess the statistical reliability of the observed performance differences in depth, we applied a paired bootstrap resampling procedure (2,000 iterations) to compute 95% confidence intervals for MUE for each method, and to evaluate the significance of all pairwise MUE differences (**Figure 6**). The bootstrap confidence intervals for MUE span approximately 0.3∼0.4 kcal/mol for all four methods, reflecting the inherent variability in estimating accuracy from 80 transformations. GROMACS-NETI yields the lowest point-estimate MUE (0.99 kcal/mol; 95% CI [0.80, 1.20]), followed by Amber-TI (1.04 kcal/mol; [0.82, 1.26]), BLaDE-MSLD (1.07 kcal/mol; [0.87, 1.29]), and OpenMM-FEP (1.11 kcal/mol; [0.88, 1.36]). The confidence intervals for all four methods overlap substantially, indicating that the observed ranking should be interpreted with caution.

**Figure 6.**
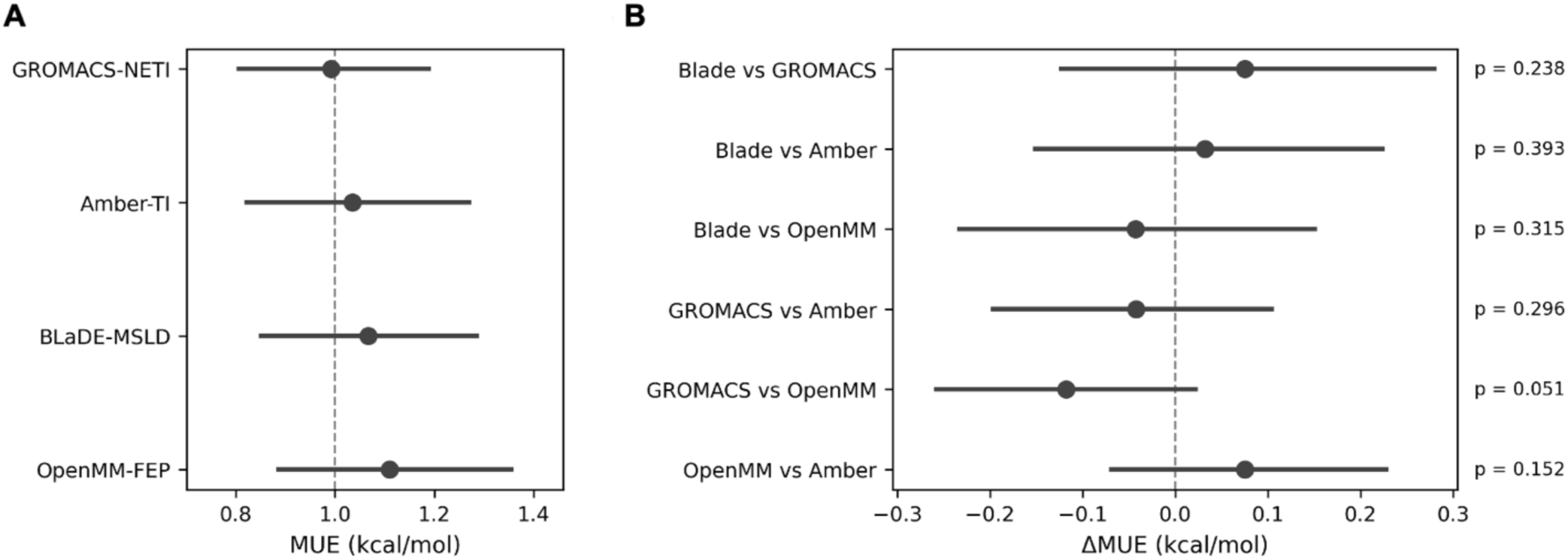
Bootstrap confidence intervals (95% CI, 2,000 iterations) for AFES method performance. (A) Point estimates and 95% CIs for MUE (kcal/mol) for each method. (B) Pairwise ΔMUE comparisons with 95% CIs and one-sided *p*-values. A vertical dashed line at zero indicates no difference.

Pairwise bootstrap comparisons confirm that no method differs significantly from any other at the conventional 0.05 level (one-sided; **Figure 6B**). GROMACS-NETI versus OpenMM-FEP has the most significant comparison (ΔMUE = −0.12 kcal/mol; 95% CI: [-0.25, +0.03]; *p* = 0.05), suggesting a modest trend favoring GROMACS-NETI that is not yet strongly supported by the current dataset size. All other pairwise differences are consistent with chance variation (*p* > 0.15). These results indicate that, while GROMACS-NETI numerically outperforms the other approaches in the JACS benchmark, the four methods are statistically equivalent at the level of the present dataset, and no definitive conclusion about superiority can be drawn from these 80 transformations alone.

#### 3.1.4 Effect of Combining Multiple Methods

Given the availability of predictions from four independent simulation approaches, we evaluated whether an ensemble of methods could provide more accurate predictions than any individual method alone (**Figure 7**). We compared three ensemble strategies; i) equal-weight averaging of all four predictions, ii) inverse-MUE-weighted averaging, and iii) an idealized oracle that selects the best-performing method for each protein. As an additional reference, we also tested iv) a linear stacking model trained via leave-one-protein-out cross-validation (LOPO-CV).

**Figure 7.**
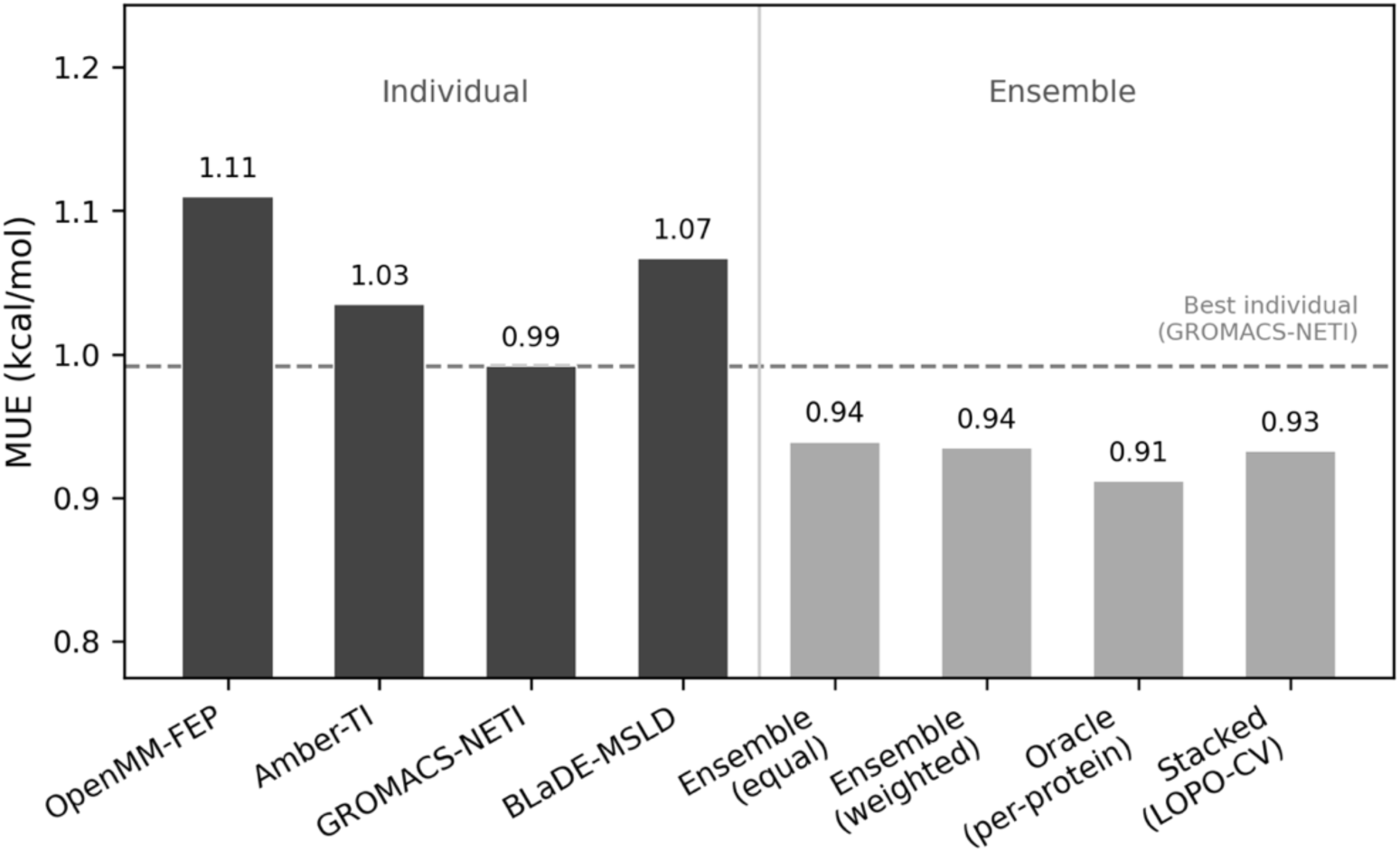
MUE for individual AFES methods and ensemble combination strategies. The dashed line indicates the MUE of the best individual method (GROMACS-NETI, 0.99 kcal/mol). Ensemble strategies include equal-weight averaging, inverse-MUE-weighted averaging, an idealized oracle (best method per protein), and a linear stacking model (LOPO-CV).

Equal-weight ensemble averaging reduced MUE from 0.99 kcal/mol (best individual, GROMACS-NETI) to 0.94 kcal/mol, a modest improvement of approximately 5%. Inverse-MUE weighting yielded a nearly identical result (0.94 kcal/mol), and even the oracle, which has access to protein-level ground truth for method selection, achieved only 0.91 kcal/mol, suggesting a ceiling on the benefit achievable by combining these four methods. The linear stacking model shows comparable MUE (0.93 kcal/mol).

The limited benefit of ensemble combination can be attributed to the strong inter-method correlation of the four predictions: Pearson correlation coefficient ranges from 0.61 (GROMACS-NETI vs. BLaDE-MSLD) to 0.84 (OpenMM-FEP vs. Amber-TI), indicating that the four methods largely agree on the same relative binding trends (data not shown). When individual predictions are highly correlated, averaging primarily reduces noise rather than systematic bias, yielding diminishing returns. In contrast, if methods with greater diversity were combined (e.g., those using different force fields or sampling protocols), the benefit of such an ensemble approach would likely be more pronounced. Exploring such heterogeneous ensembles represents an interesting direction for future work.

#### 3.1.5 Effect of Replica Averaging

Each of the four methods was run with four independent replicas. To evaluate the benefit of averaging multiple replicas, we systematically computed the mean MUE across all combinations of k replicas for k = 1, 2, 3, 4 (**Figure 8**). This analysis quantifies how much accuracy can be gained by running additional replicates per transformation.

**Figure 8.**
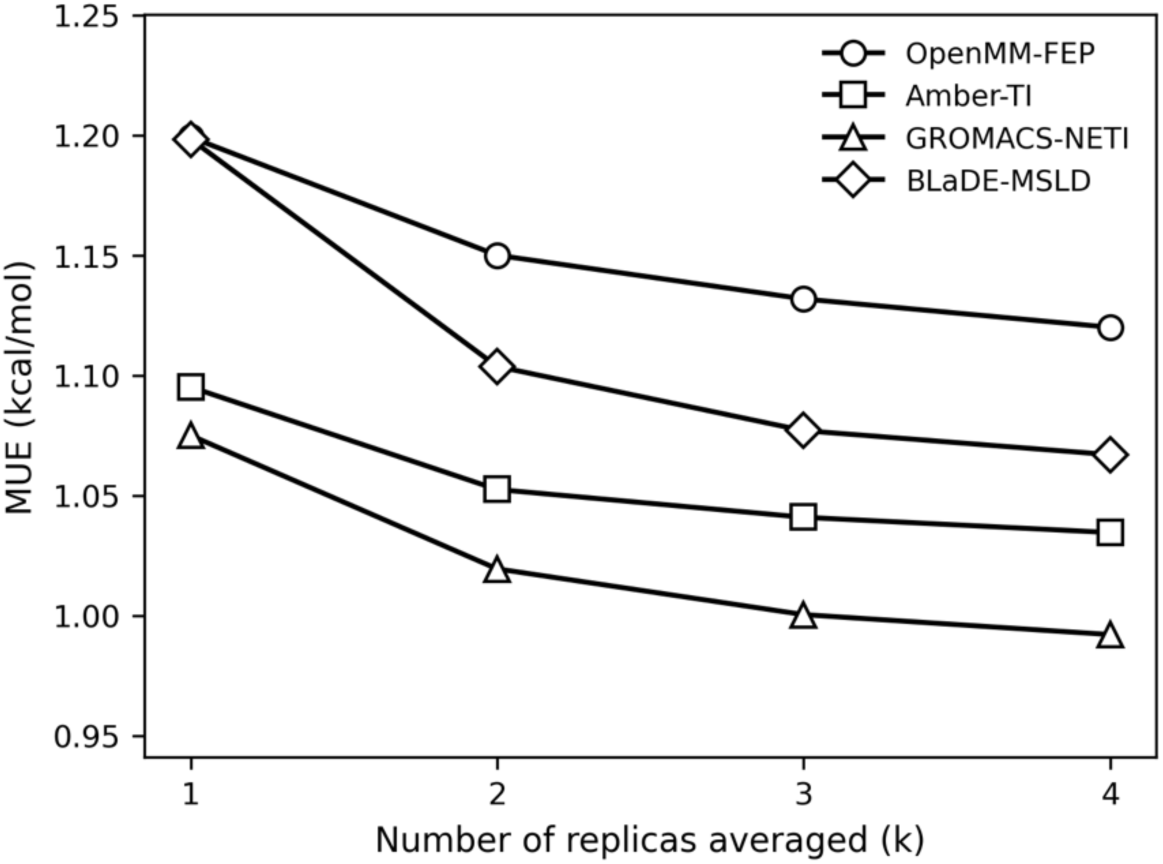
MUE as a function of the number of replicas averaged (k = 1, 2, 3, 4) for each of the four AFES methods, computed over all C(4, k) replica combinations.

For all four methods, averaging replicas reduces MUE relative to a single replica, but the gains are modest. Going from one to four replicas improves MUE by 6∼11% across methods: BLaDE-MSLD shows the largest absolute gain (1.20 → 1.07 kcal/mol; 11%), followed by GROMACS-NETI (1.08 → 0.99 kcal/mol; 7.7%), OpenMM-FEP (1.19 → 1.11 kcal/mol; 6.6%), and Amber-TI (1.10 → 1.04 kcal/mol; 5.5%). The most substantial fraction of the total gain occurs in the first additional replica; the 1 → 2 step accounts for 50∼73% of the full 1 → 4 improvement across methods. In other words, beyond k = 2, the marginal benefit diminishes rapidly. These results suggest that a minimum of two replicas is advisable to capture the most impactful variance reduction, while the incremental benefit of running three or four replicas is relatively small given the additional computational cost.

#### 3.1.6 Computational Performance

To compare the computational performance of each AFES method, we first evaluated ten thrombin morphs (i.e., transformations) using Amber-TI, GROMACS-NETI, and OpenMM-FEP on an NVIDIA RTX A5000 GPU (**Table 2** and **Table S6**). The total wall-clock time is 135.51 h (13.55 h per morph) for Amber-TI, 118.34 h (11.83 h per morph) for GROMACS-NETI, and 122.08 h (12.21 h per morph) for OpenMM-FEP, indicating that the nonequilibrium protocol is approximately 9∼13% faster on average in our current simulation protocol. For every individual morph, the total runtime of GROMACS-NETI was shorter than that of Amber-TI and OpenMM-FEP, and the performance advantage was particularly pronounced for the ligand leg of the calculation (**Table S6**).

**Table 2.**
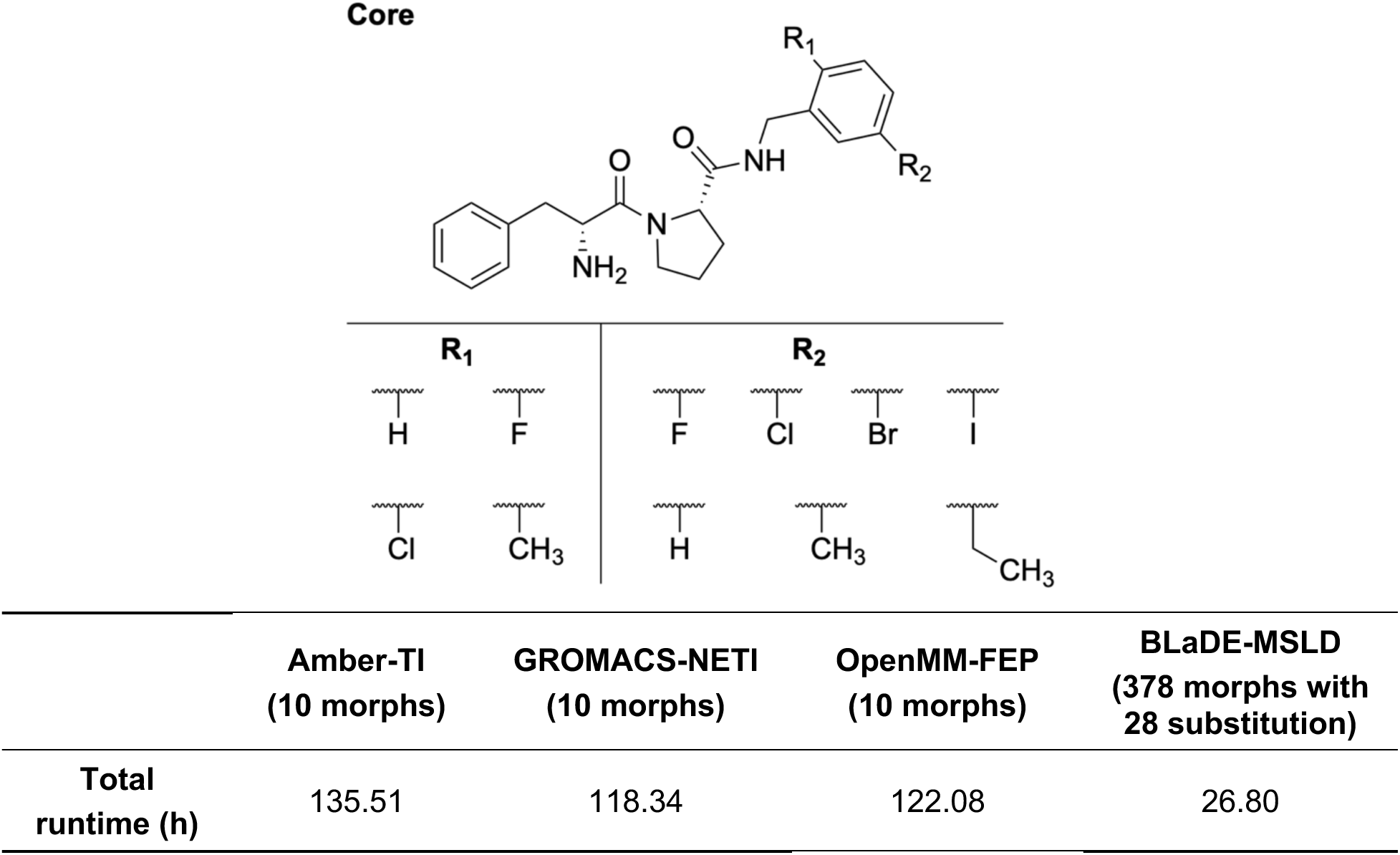
Performance comparison of four AFES methods for thrombin using one NVIDIA RTX A5000 GPU.

In BLaDE-MSLD, a single multi-ligand simulation samples all 28 substitution combinations (4 from site R1 × 7 from site R2 in **Table 2**), and the relative free energies of the individual ligands are extracted from this trajectory. These per-ligand free energies can be combined to construct a total of 378 morph sets, corresponding to all possible unordered ligand pairs, without performing separate simulations for each pair. In other words, the 378 morphs are obtained by post-processing a single simulation of all 28 substitution combinations from 10 ligands rather than 378 distinct alchemical transformations. The total wall-clock time required for this BLaDE-MSLD simulation is 26.80 h with 2-fs timestep (n.b., GROMACS-NETI and Amber-TI used 4-fs timestep with HMR in **Table 2**).

For the TI protocols and the FEP calculations, the mean total runtime averaged over 80 transformations in Amber-TI, GROMACS-NETI, and OpenMM-FEP is 12.53 h per morph. In contrast, BLaDE-MSLD characterized 378 morph sets in 26.80 h, which corresponds to an effective cost of approximately 0.07 h per morph. These results demonstrate that the MSLD approach provides substantially higher computational efficiency and throughput in terms of the number of ligand pairs that can be characterized per unit wall-clock time. A caveat is that Amber-TI, GROMACS-NETI, and OpenMM-FEP calculations could easily be distributed over many GPUs simultaneously, effectively lowering wall-clock time (e.g., using 12 GPUs, Amber-TI, GROMACS-NETI, and OpenMM-FEP calculations could be done a little more than 1 hour). Such a wall-clock time reduction cannot be done easily with BLaDE-MSLD.

### 3.2 Case Study 1: Glucocorticoid Receptor

To demonstrate the practical application of *MolCube-AFES*, we conducted a case study using the glucocorticoid receptor (GR, PDB ID 3BQD), which was co-crystallized with deacylcortivazol.^48^ The 3BQD structure was originally used in the study by *Weinstein et al.* to model mode of azaxanthene-based selective GR modulators.^16^ This system serves as a benchmark due to the availability of a comprehensive experimental dataset, comprising 13 compounds in total, including compound 14 as the reference, which cover a range of binding affinity (Ki values from 1.0 to 49.5 nM) reflecting modest energetic variations across various substituents.

In this study, we selected compound 14 from this series as the starting reference ligand because it possesses the most basic structure with an unsubstituted phenyl ring (R=H) (**Table S7**). However, because the protein-ligand complex structure with compound 14 is unavailable and the deacylcortivazol scaffold differs significantly from the azaxanthene core, identifying an appropriate binding mode is essential. Specifically, deacylcortivazol consists of five fused rings with a negative surface potential, whereas compound 14 features a three-ring core with a sulfur-containing thiazole and a more positive charge distribution. Due to these fundamental structural differences, we utilized Boltz-2^26^ to generate the binding pose for the receptor-compound 14 complex. Subsequent analysis of the generated structure reveals a distinct binding mode (**Figure 9**). While the interaction with GLN-570 was preserved, the polar interactions with ASN-564 and GLN-642 observed in 3BQD were lost as these residues encountered the hydrophobic regions of compound 14. Instead, new critical interaction was established with ASN-611. The predicted orientation indicates that while compound 14 occupies the same binding site as deacylcortivazol, its unique scaffold leads to a reorganized interaction network within the pocket.

**Figure 9.**
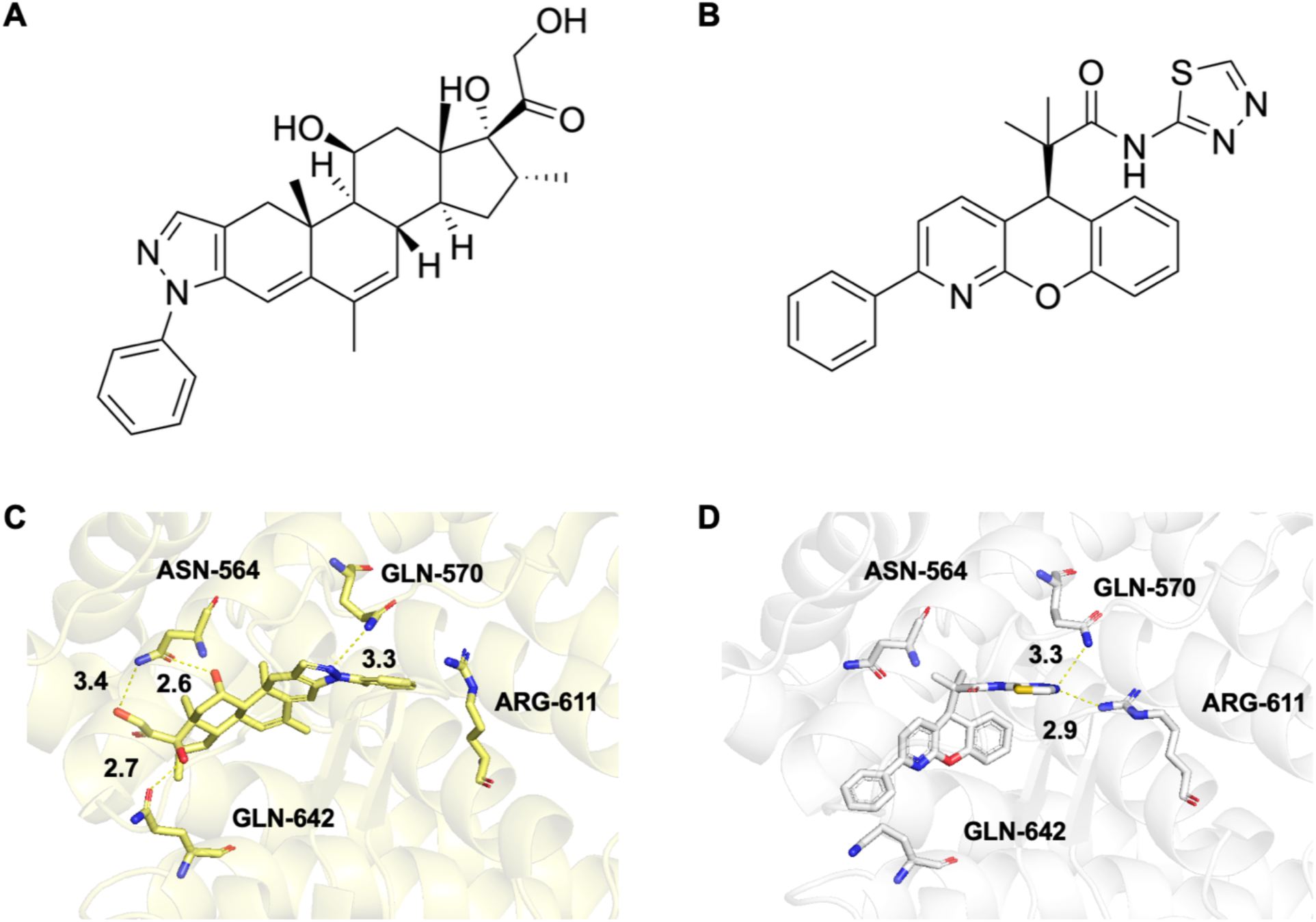
Structural comparison between the native ligand and the modeled compound. (A) 2D structure of deacylcortivazol, the co-crystallized ligand in PDB ID 3BQD. (B) 2D structure of compound 14, selected as the starting reference ligand for ΔG_bind_ calculations. (C) Detailed binding mode of deacylcortivazol in 3BQD, showing key polar interaction with residues ASN-564 and GLN-642. (D) Predicted binding mode of compound 14 generated by Boltz-2. The interaction with GLN-570 is preserved, whereas contacts with ASN-564 and GLN-642 are lost and a new critical interaction with ARG-611 is established.

A total of 13 ligands with three substitution sites were tested, featuring various substitutions including alkyl chains (methyl, ethyl, propyl), heteroatom-containing groups (methoxy, isopropoxy, dimethylamino, hydroxy), and carbonyl derivatives (acetyl, isopropanoyl) (**Table S7**). To enhance the accuracy of simulation, the ligands were grouped based on their structural similarity by considering both the substitution sites and the chemical nature of the substituents to guide perturbation selection in the Amber-TI, GROMACS-NETI and OpenMM-FEP calculations. This classification reflects gradual changes in steric bulk and polarity, providing a practical framework for consistent comparison of calculated binding free energies while maintaining numerical stability in AFES. A comprehensive alchemical perturbation network was constructed connecting compounds 14 through 28 in multiple closed cycles (**Figure S2**). Crucially, each perturbation link (e.g., 14 to 15) was carefully designed to maximize structural overlap and enhance computational efficiency by representing the smallest feasible systematic substitution of functional groups (like Me or OMe), reflecting the gradual chemical variation within the congenic series. For the BLaDE-MSLD calculations, all ligands were handled as a single group, and the same set of perturbation networks in **Figure S2** was used for the comparison with Amber-TI, GROMACS-NETI, and OpenMM-FEP calculations.

The predicted ΔΔG_bind_ are compared with the experimental data for a congeneric ligand series, showing modest energetic variation within 2 kcal/mol (**Figure 10**). All four computational methods show MUE below 1 kcal/mol, comparable to the typical experimental uncertainty of binding measurements. Notably, BLaDE-MSLD shows the most consistent performance (RMSE = 0.82 kcal/mol, MUE = 0.68 kcal/mol), indicating robust quantitative accuracy within the narrow ΔΔG_bind_ range of this dataset, whereas OpenMM-FEP yielded results comparable to other standard AFES methods (RMSE=1.07 kcal/mol, MUE = 0.98 kcal/mol). Although absolute error-based analysis shows promising agreement with experiment, the coefficients of determination (R²) are low and are not considered reliable indicators of model accuracy (**Table 3**). This is primarily attributed to the modest energetic variation of the congeneric ligand series. Taken together, the low MUE and RMSE values, despite the close-to-zero R² arising from the limited energetic dynamic range of the series, indicate that the method can reliably resolve structurally meaningful energetic differences within this congeneric ligand set.

**Figure 10.**
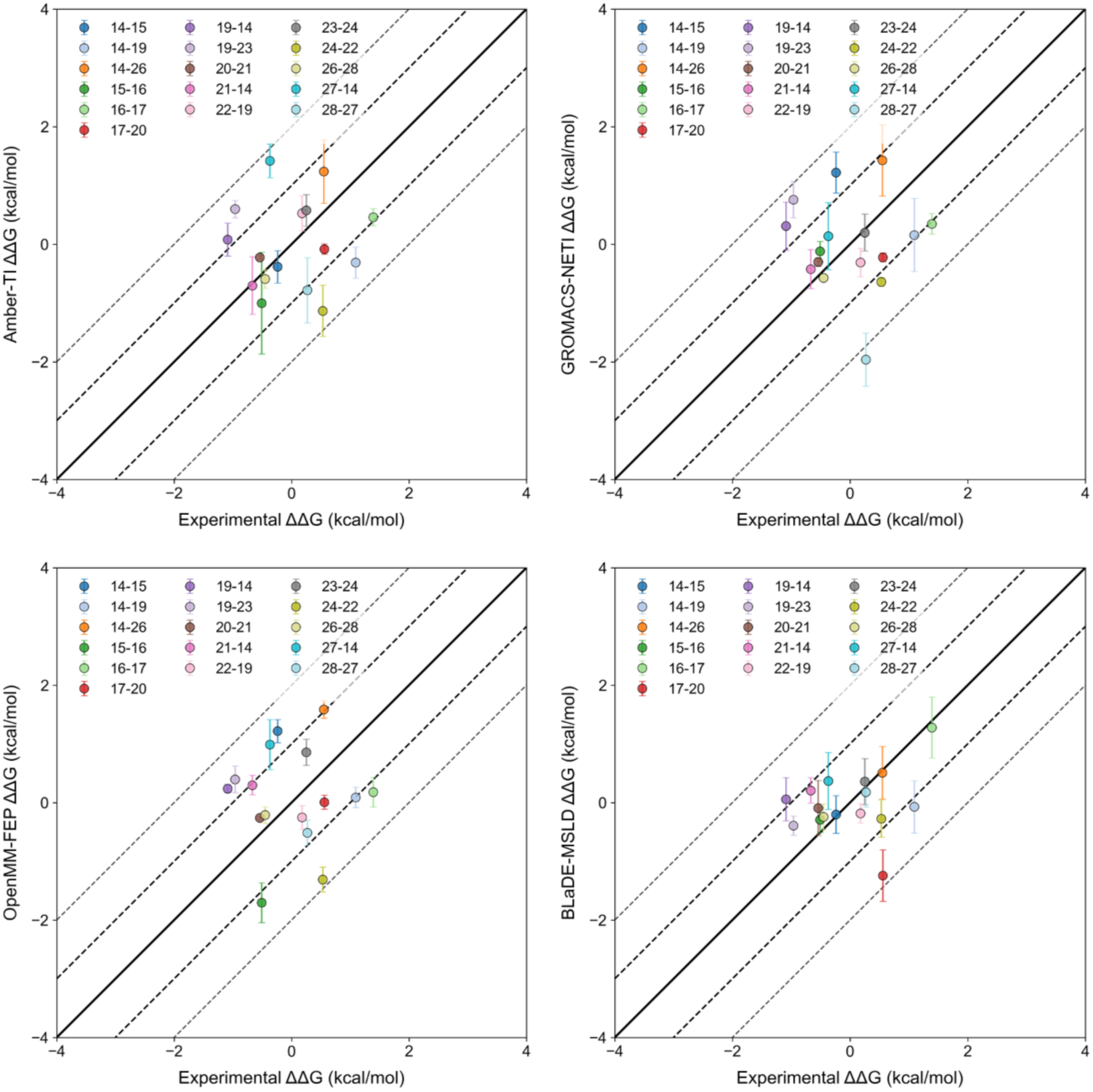
ΔΔG_bind_ prediction-experiment for the glucocorticoid receptor case study.

**Table 3.**
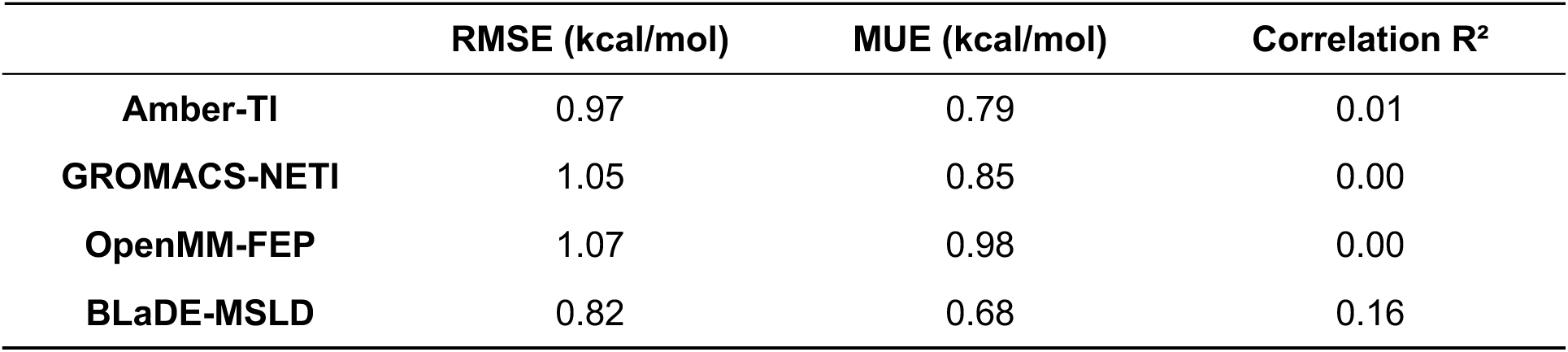
Comparison of MUE, RMSE, and R^2^ for the glucocorticoid receptor case study.

### 3.3 Case Study 2: HDAC6 inhibitor

The study by *Wang et al.*^17^ reported TNI-97 as a highly selective and orally bioavailable HDAC6 inhibitor with promising activity against triple-negative breast cancer. HDAC6 was therefore selected as the target for our AFES benchmark, as the TNI-97 series (18 compounds: 30a-30r in **Table S8**) offers a well-defined and experimentally characterized dataset with reported IC₅₀ values suitable for rigorous computational evaluation. For the structural basis of the simulations, we employed the docking structure of HDAC6 with compound 30a (based on PDB ID 5EDU)^49^ provided by the original authors (**Figure 11**). All ligands in the series share a conserved 5-pyrazolyl-1H-benzotriazole scaffold appended with a terminal hydroxamate group responsible for zinc coordination, while systematic substitutions around the benzotriazole and phenyl moieties were introduced to modulate isoform (HDAC1) selectivity (**Table S8**). To ensure robust ΔΔG_bind_ calculations, we constructed an alchemical perturbation network connecting compounds 30a through 30r in a closed cycle, enabling internal consistency (**Figure S3**).

**Figure 11.**
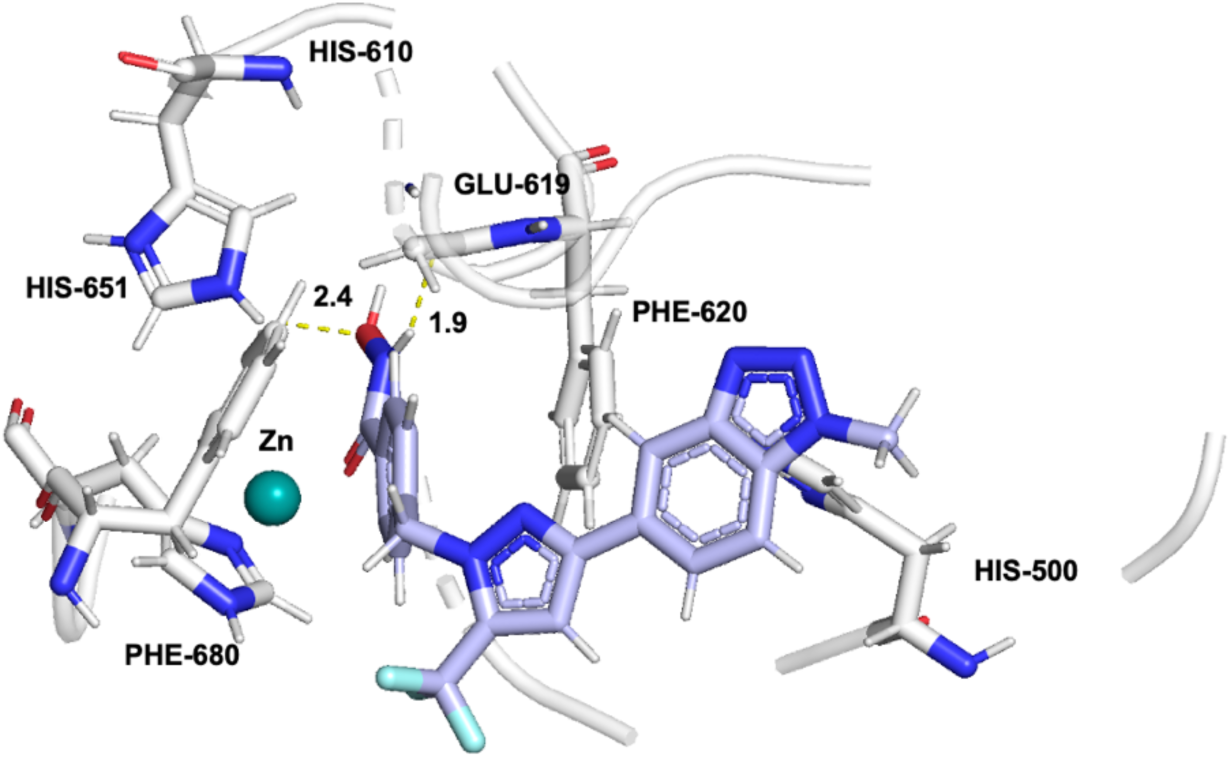
Docking structure of HDAC6 (5EDU) with compound 30a.

The predicted ΔΔG_bind_ from all four methods exhibit an MUE approaching 2 kcal/mol (**Figure 12** and **Table 4**). While this level of deviation necessitates caution regarding absolute quantitative prediction, the MUE is commensurate with the typical uncertainty of binding assays. All four methods reproduced similar qualitative ranking trends for the subset of ligands, although global correlations were limited by specific outliers. Consequently, the global R^2^ values are close to 0 for all methods, reflecting substantial deviations from experimental values in a subset of morphs involving a specific ligand, rather than a systematic deficiency of the methods themselves.

**Figure 12.**
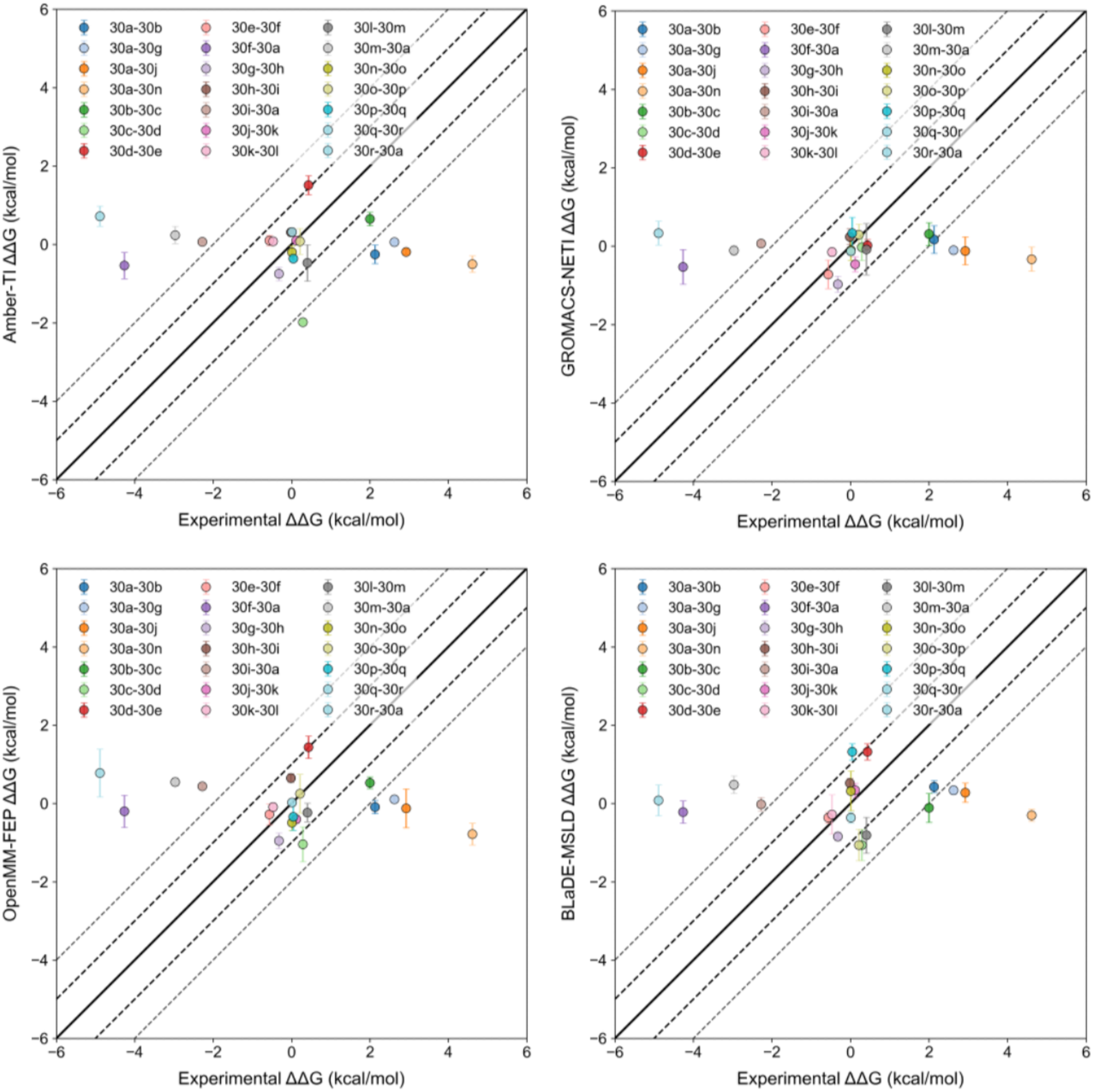
ΔΔG_bind_ prediction-experiment for the HDAC6 inhibitor case study.

**Table 4.** Comparison of MUE, RMSE, and R^2^ for the HDAC6 inhibitor case study.

| | RMSE (kcal/mol) | MUE (kcal/mol) | Correlation $R^2$ |
| --- | --- | --- | --- |
| Amber-TI | 2.44 | 1.76 | 0.08 |
| **GROMACS-NETI** | 2.23 | 1.54 | 0.00 |
| **OpenMM-FEP** | 2.38 | 1.75 | 0.02 |
| **BLaDE-MSLD** | 2.29 | 1.75 | 0.00 |

Transformations involving 30a exhibit the most significant deviations from the experimental values (see the raw data in **AFES_Supplementary2.xlsx**). For instance, in the case of 30a to 30b transformation, the computed ΔΔG_bind_ was nearly zero, a result physically consistent with the high solvent accessibility of the perturbation site. Despite this, the experimental data show a pronounced affinity difference (2.13 kcal/mol). This discrepancy suggests that the experimental trend may not arise from direct protein-ligand interactions but could instead be an artifact of binding mode instability or specific assay conditions unique to 30a. While simulations reached adequate convergence, the stark contrast with experimental data suggests that the latter may reflect complex structural transitions or solvent effects that challenge the standard interpretation of binding affinity at this position.^50^

To quantitatively isolate the source of this inconsistency, we converted the relative binding free energies (ΔΔG_bind_) into the absolute binding free energies (ΔG_bind_) to allow direct comparison with experimental data across multiple simulations. All absolute free energies were reconstructed from the relative free energy network by solving an overdetermined system of linear equations using a least-squares approach with a zero-mean gauge constraint. In addition, a single global offset was applied for comparison with the experimental data. Consequently, converting ΔΔG_bind_ to ΔG_bind_ led to a significant improvement in predictive accuracy, and 30a becomes more clearly identifiable as an outlier (see the raw data in **AFES_Supplementary2.xlsx**). Upon conversion to ΔG_bind_, the error metrics improved to RMSE/MUE values of 1.32/1.02 kcal/mol for GROMACS-NETI, followed by Amber-TI (RMSE = 1.56 kcal/mol, MUE = 1.33 kcal/mol), OpenMM-FEP (RMSE = 1.51 kcal/mol, MUE = 1.20 kcal/mol), and BLaDE-MSLD (RMSE = 1.46 kcal/mol, MUE = 1.25 kcal/mol) (**Figure 13**). These reduced error metrics confirm that the earlier discrepancies in ΔΔG_bind_ were largely artifacts of error propagation from the 30a outlier rather than a systematic deficiency of the AFES methods themselves.

**Figure 13.**
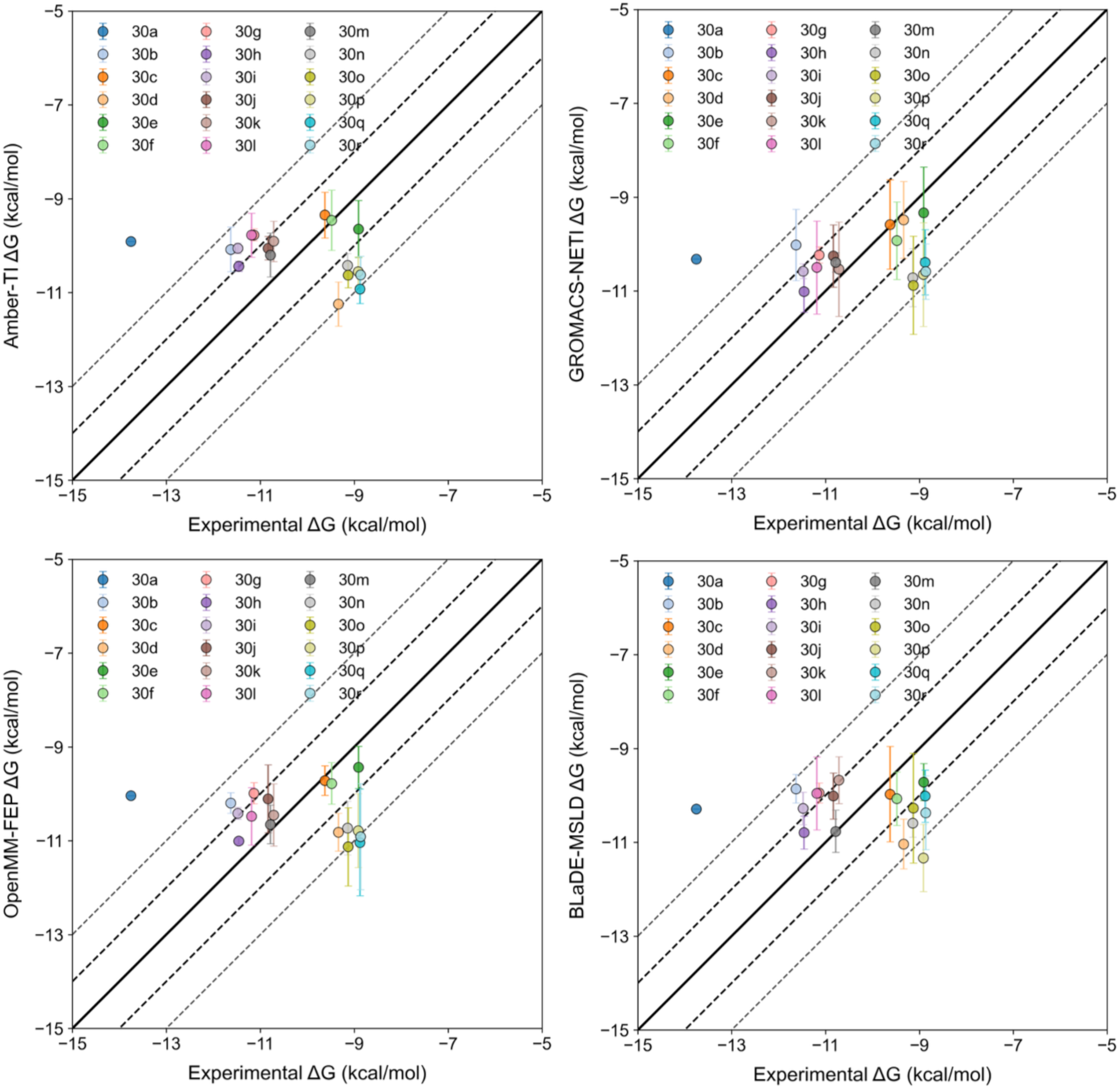
Absolute binding free energy (ΔG_bind_) prediction-experiment for the HDAC6 inhibitor case study.

## 4. Conclusions

The comparative evaluation of Amber-TI, GROMACS-NETI, OpenMM-FEP, and BLaDE-MSLD for the JACS benchmark set and 2 case studies (GR and HDAC6) has revealed broadly comparable performance profiles, with quantitative differences that are modest in magnitude. GROMACS-NETI achieved the lowest MUE (0.99 kcal/mol), closely followed by Amber-TI (MUE = 1.03 kcal/mol), BLaDE-MSLD (MUE = 1.07 kcal/mol), and OpenMM-FEP (MUE = 1.11 kcal/mol). Importantly, bootstrap analysis (2,000 iterations) demonstrates that the 95% confidence intervals for all four methods overlap substantially, and pairwise differences are not statistically significant (*p* > 0.05 for all comparisons). These findings indicate that the four methods are statistically equivalent in the present benchmark and that observed ranking differences reflect sampling variability rather than systematic performance differences. Across all methods, predictions capture experimental trends within the margin of error that support actionable interpretation in a drug-discovery context.

Given the statistical equivalence of the four methods, practical method selection should be guided by workflow requirements rather than expected accuracy differences. GROMACS-NETI and Amber-TI are well suited to campaigns where absolute accuracy and robust rank ordering are the primary decision metric. In our testing, GROMACS-NETI shows a modest numerical edge against other methods. OpenMM-FEP offers broad compatibility through its open-source ecosystem and is appropriate when portability and extensibility are priorities. BLaDE-MSLD remains the method of choice for high throughput evaluation of extensive ligand sets because MSLD enables simultaneous sampling of many candidates in a single simulation, improving turnaround without sacrificing the quality required for screening. Therefore, method selection should align with the decision metric of interest, the anticipated magnitude of binding affinity changes, and the computational constraints of the project.

Ensemble averaging across the four methods yields only a modest improvement. About 7% MUE is reduced relative to the best individual method. This is consistent with the high inter-method correlation observed throughout the benchmark. This limited gain implies that the four approaches share common systematic error sources (most likely force field parameters and perturbation network design). Therefore, combining predictions does not substantially cancel out errors. Similarly, replica averaging exhibits strong diminishing returns. Increasing from one to two replicas captures 50∼73% of the total improvement achievable by averaging all four replicas, while additional replicas contribute progressively less. Together, these results suggest that the dominant error source in current AFES workflow is systematic rather than statistical, and that investment in the force field quality, perturbation-network design, and structural input quality are likely to yield greater accuracy improvements than simply increasing replica count or combining methods.

The scope of this evaluation is restricted to 10 protein targets (8 from the JACS set and 2 case studies) and a relatively small set of ligand transformations, which limits the generalizability of the findings to broader chemical spaces. Each method also exhibits inherent sampling constraints that may affect convergence for complex transformations. TI relies on sufficient overlap between λ-windows, nonequilibrium switching in NETI introduces sensitivity to transition protocols, and MSLD requires careful parameterization and/or grouping to ensure stability and accuracy. Protein-ligand complex structural biases represent another source of uncertainty, as docking-derived initial poses and softcore parameter settings can influence the accuracy of predicted free energy changes, particularly for transformations involving ambiguous binding modes. Furthermore, the experimental datasets used in the two case studies exhibit a narrow range of ΔΔG_bind_ values, which reduces the interpretability of correlation metrics as indicators of predictive performance and may obscure systematic errors. These limitations highlight the need for caution when extrapolating the results to systems with greater structural diversity or more pronounced energetic differences.

Future work should explore hybrid approaches that combine the strengths of these methods, such as integrating ligand grouping strategies for MSLD to improve sampling efficiency. Based on our internal testing, split-group configurations generally yield higher accuracy than single-group projects. However, aside from unavoidable edge cases such as p38 and PTP1B, these improvements are typically modest and, in some instances, may slightly decrease accuracy. To ensure higher numerical robustness, we recommend running at least 2 independent replicas to reduce statistical noise and enhance confidence in the final estimates. *MolCube-AFES* supports automatic ligand grouping for BLaDE-MSLD system generation, with the maximum number of substituents per site as 10 to ensure robust and broadly applicable performance.

Separately from sampling considerations, the availability of reliable structural priors is a critical factor for achieving accurate free energy predictions, particularly for complex protein-ligand systems. Recent advances in AI-based algorithms for protein-ligand complex structure generation^51^ provide a powerful means of supplying high-quality initial structures for downstream free energy calculations. The effectiveness of this approach is illustrated by our GR case study, in which protein-ligand complex structures generated using Boltz-2 were subsequently used to initialize AFES calculations, yielding high quantitative accuracy (MUE of around 1 kcal/mol) despite the narrow ΔΔG_bind_ range of the dataset. These results highlight the potential of integrating AI-derived structural models with physics-based free energy methods. Looking forward, further coupling such structure generation approaches with GPU-optimized free energy algorithms offers a promising pathway to improving robustness, accuracy, and scalability in drug discovery applications.

## Supporting information

Supporting Raw Data

## Acknowledgements

This work is supported by the Scale-up TIPS Program (RS-2023-00321786) funded by the Ministry of SMEs and Startups (MSS, Korea).

## Conflict of Interest

W.I. is the co-founder and CEO of MolCube INC.

## Data Availability

All input files used in this study are available at https://github.com/molcube/molcube_afes_rbfe.

