## Supporting Tables and Figures for "Comparative Analysis of Relative Ligand Binding Free Energy Simulation Methods: Amber-TI, GROMACS-NETI, OpenMM-FEP, and BLaDE-MSLD"

**Table S1.** 10 transformations of 8 proteins in the JACS set.

| Protein | RCSB ID | Ligand 1 | Ligand 2 |
|---|---|---|---|
| BACE | 4DJW | CAT-4k | CAT-4o |
| | | CAT-13o | CAT-17h |
| | | CAT-17b | CAT-17e |
| | | CAT-24 | CAT-17i |
| | | CAT-4m | CAT-4p |
| | | CAT-4m | CAT-13j |
| | | CAT-13d | CAT-13b |
| | | CAT-13d | CAT-13h |
| | | CAT-13g | CAT-17i |
| | | CAT-13j | CAT-4o |
| Tyk2 | 4GIH | jmc_23 | ejm_46 |
| | | jmc_23 | ejm_55 |
| | | jmc_23 | jmc_30 |
| | | ejm_47 | ejm_31 |
| | | ejm_49 | ejm_31 |
| | | ejm_55 | ejm_54 |
| | | ejm_31 | ejm_48 |
| | | ejm_31 | ejm_45 |
| | | ejm_43 | ejm_55 |
| | | ejm_44 | ejm_42 |
| CDK2 | 1H1Q | 22 | 1h1r |
| | | 26 | 1oi9 |
| | | 30 | 26 |
| | | 31 | 32 |
| | | 1h1r | 1oi9 |
| | | 1h1s | 26 |
| | | 1oi9 | 20 |
| | | 1oiy | 1h1q |
| | | 17 | 1h1q |
| | | 20 | 1h1q |
| MCL1 | 4HW3 | 35 | 33 |
| | | 38 | 35 |
| | | 42 | 51 |
| | | 43 | 47 |
| | | 26 | 57 |
| | | 62 | 45 |
| | | 28 | 35 |
| | | 67 | 35 |
| | | 67 | 53 |
| | | 67 | 27 |

| | | | |
|---|---|---|---|
| JNK1 | 2GMX | 18634-1 | 18637-1 |
| | | 18636-1 | 18624-1 |
| | | 18637-1 | 18631-1 |
| | | 18638-1 | 18634-1 |
| | | 18659-1 | 18634-1 |
| | | 18626-1 | 18634-1 |
| | | 18626-1 | 18658-1 |
| | | 18626-1 | 18659-1 |
| | | 18626-1 | 18627-1 |
| | | 18626-1 | 18628-1 |
| p38 | 3FLY | p38a_2l | p38a_2j |
| | | p38a_2l | p38a_2o |
| | | p38a_2m | p38a_2j |
| | | p38a_2m | p38a_2k |
| | | p38a_2s | p38a_2l |
| | | p38a_2v | p38a_3fln |
| | | p38a_2v | p38a_3fmk |
| | | p38a_3fly | p38a_2x |
| | | p38a_3fly | p38a_3flq |
| | | p38a_3flz | p38a_2c |
| thrombin | 2ZFF | 1a | 5 |
| | | 6e | 6b |
| | | 1b | 1a |
| | | 1b | 3b |
| | | 1d | 1a |
| | | 1d | 1c |
| | | 1d | 5 |
| | | 3a | 1b |
| | | 6a | 1b |
| | | 6a | 6b |
| PTP1B | 2QBS | 20667(2qbp) | 23479 |
| | | 23471 | 23468 |
| | | 23477 | 23467 |
| | | 23482 | 23479 |
| | | 23484 | 23486 |
| | | 20670(2qbs) | 23477 |
| | | 20670(2qbs) | 23466 |
| | | 23467 | 23475 |
| | | 23467 | 23476 |
| | | 23467 | 23466 |

**Table S2.** Comparison of $\Delta\Delta G_{bind}$ values across the four methods with experimental results.

| Protein | Amber-TI | | | | GROMACS-NETI | | | | OpenMM-FEP | | | | BLaDE-MSLD | | | |
|---|---|---|---|---|---|---|---|---|---|---|---|---|---|---|---|---|
| | RMSE[a] | MUE[a] | $R^2$ | $r_P$ | RMSE[a] | MUE[a] | $R^2$ | $r_P$ | RMSE[a] | MUE[a] | $R^2$ | $r_P$ | RMSE[a] | MUE[a] | $R^2$ | $r_P$ |
| **BACE** | 1.16 | 0.88 | 0.46 | 0.68 | 1.12 | 0.86 | 0.45 | 0.67 | 1.14 | 0.98 | 0.14 | 0.37 | 0.88 | 0.61 | 0.60 | 0.78 |
| **Tyk2** | 1.05 | 0.81 | 0.51 | 0.71 | 1.09 | 0.85 | 0.42 | 0.64 | 1.14 | 0.85 | 0.34 | 0.58 | 1.05 | 0.89 | 0.54 | 0.73 |
| **CDK2** | 1.13 | 0.79 | 0.44 | 0.66 | 1.04 | 0.75 | 0.48 | 0.69 | 1.07 | 0.84 | 0.47 | 0.68 | 1.27 | 0.96 | 0.28 | 0.53 |
| **MCL1** | 2.30 | 1.72 | 0.26 | 0.51 | 2.04 | 1.64 | 0.29 | 0.54 | 2.48 | 1.84 | 0.27 | 0.52 | 2.37 | 1.99 | 0.16 | 0.40 |
| **JNK1** | 1.15 | 0.97 | 0.26 | 0.51 | 1.30 | 1.04 | 0.18 | 0.43 | 1.32 | 1.13 | 0.14 | 0.38 | 1.07 | 0.80 | 0.11 | -0.33 |
| **p38** | 1.58 | 1.23 | 0.51 | 0.71 | 1.15 | 0.99 | 0.66 | 0.81 | 1.45 | 1.08 | 0.53 | 0.73 | 1.06 | 1.30 | 0.43 | 0.65 |
| **thrombin** | 0.61 | 0.45 | 0.36 | 0.60 | 0.73 | 0.56 | 0.39 | 0.63 | 0.62 | 0.50 | 0.48 | 0.70 | 0.98 | 0.73 | 0.39 | 0.63 |
| **PTP1B** | 2.02 | 1.43 | 0.01 | 0.12 | 1.76 | 1.25 | 0.20 | 0.45 | 2.32 | 1.62 | 0.01 | -0.10 | 1.84 | 1.26 | 0.13 | 0.36 |

[a]The unit is kcal/mol.

**Table S3.** Systems used in BLaDE-MSLD. The group id, number of ligands, number of sites, and number of substituents are listed.

| Protein | Group ID | # ligands | # Sites | # Subs |
|---|---|---|---|---|
| BACE | | 15 | 2 | 7 × 9 |
| Tyk2 | | 13 | 1 | 13 |
| CDK2 | | 12 | 2 | 6 × 7 |
| MCL1 | | 15 | 2 | 12 × 7 |
| JNK1 | | 11 | 4 | 3 × 4 × 4 × 4 |
| p38 | p38-1 | 14 | 3 | 5 × 6 × 6 |
| | p38-2a | 6 | 1 | 6 |
| | p38-2b | 4 | 1 | 4 |
| | p38-2c | 6 | 1 | 6 |
| thrombin | | 10 | 2 | 4 × 7 |
| PTP1B | PTP1B-1 | 13 | 2 | 3 × 11 |
| | PTP1B-2a | 5 | 2 | 3 × 3 |
| | PTP1B-2b | 9 | 1 | 9 |

**Table S4.** $\Delta\Delta G_{bind}$ prediction difference in BLaDE-MSLD using 1 group and split groups.

| Protein | Ligand 1 | Ligand 2 | $\Delta\Delta G_{1group}$ | $\Delta\Delta G_{groups}$ | exp. |
|---|---|---|---|---|---|
| p38 | p38a_2l | p38a_2j | -1.82 | -1.34 | 2.18 |
| | p38a_2s | p38a_2l | 0.28 | -0.76 | -1.15 |
| | p38a_3fly | p38a_2x | 0.13 | 0.23 | 1.19 |
| | p38a_3flz | p38a_2c | 0.06 | -1.84 | -0.97 |
| | p38a_2l | p38a_2o | 0.76 | 1.08 | 1.77 |
| | p38a_2m | p38a_2k | 1.78 | 2.84 | 0.41 |
| | p38a_3fly | p38a_3flq | 0.13 | 0.34 | 1.49 |
| | p38a_2m | p38a_2j | -1.51 | -0.70 | 0.88 |
| | p38a_2v | p38a_3fln | -1.49 | -2.65 | -1.91 |
| | p38a_2v | p38a_3fmk | -1.54 | -3.55 | -2.86 |
| | MUE[a] | | 1.54 | 1.30 | |
| | $r_p$ | | 0.23 | 0.65 | |
| PTP1B | 23467 | 23476 | -6.30 | -1.07 | -2.07 |
| | 23471 | 23468 | -5.50 | 0.56 | 0 |
| | 20670(2qbs) | 23477 | -1.06 | 0.12 | 0.23 |
| | 23477 | 23467 | 7.09 | 0.75 | 1.51 |
| | 23467 | 23466 | 4.63 | 0.32 | -0.51 |
| | 23482 | 23479 | 33.19 | -0.51 | 0.86 |
| | 20670(2qbs) | 23466 | 10.66 | 1.19 | 1.24 |
| | 23467 | 23475 | -6.21 | -0.82 | -1.38 |
| | 23484 | 23486 | 1.08 | 0.03 | -4.72 |
| | 20667(2qbp) | 23479 | -45.28 | -0.27 | 2.28 |
| | MUE[a] | | 12.17 | 1.26 | |
| | $r_p$ | | -0.11 | 0.36 | |

[a]The unit is kcal/mol.

**Table S5.** Comparison of $\Delta\Delta G_{bind}$ values from the Wang et al. with experimental results.

| | RMSE[a] | MUE[a] | Correlation $R^2$ | $r_P$ | Acc ±1 | Acc ±2 |
|---|---|---|---|---|---|---|
| **Wang et al.** | 1.27 | 0.99 | 0.35 | 0.65 | 0.58 | 0.89 |

[a]The unit is kcal/mol.

**Table S6.** Single-GPU runtime (hours) per morph for 10 thrombin alchemical transformations.

| | Amber-TI | | | GROMACS-NETI | | | OpenMM-FEP | | |
|---|---|---|---|---|---|---|---|---|---|
| | Complex | Ligand | Total | Complex | Ligand | Total | Complex | Ligand | Total |
| 1a-5 | 9.26 | 4.22 | 13.48 | 9.33 | 3.46 | 12.78 | 10.63 | 2.73 | 13.36 |
| 6e-6b | 9.41 | 4.06 | 13.47 | 9.20 | 3.24 | 12.45 | 11.12 | 3.08 | 14.20 |
| 1b-1a | 9.23 | 4.07 | 13.30 | 8.63 | 2.10 | 10.74 | 10.43 | 2.73 | 13.16 |
| 1b-3b | 9.44 | 4.18 | 13.62 | 9.48 | 3.52 | 13.00 | 10.78 | 2.92 | 13.70 |
| 1d-1a | 9.23 | 4.33 | 13.55 | 9.59 | 3.05 | 12.64 | 10.78 | 2.75 | 13.53 |
| 1d-1c | 9.40 | 4.08 | 13.47 | 8.65 | 2.02 | 10.67 | 10.60 | 2.77 | 13.37 |
| 1d-5 | 9.11 | 4.35 | 13.45 | 9.02 | 3.10 | 12.13 | 10.58 | 2.72 | 13.30 |
| 3a-1b | 9.63 | 4.22 | 13.86 | 9.22 | 2.74 | 11.96 | 10.45 | 2.98 | 13.43 |
| 6a-1b | 9.53 | 4.17 | 13.71 | 8.45 | 2.17 | 10.63 | 10.53 | 2.93 | 13.46 |
| 6a-6b | 9.43 | 4.17 | 13.6 | 8.70 | 2.63 | 11.34 | 10.88 | 3.05 | 13.93 |
| Avg. | | | 13.55 | | | 11.83 | | | 13.54 |

**Table S7.** Chemical structures of azaxanthene-based compounds used for AFES calculations.

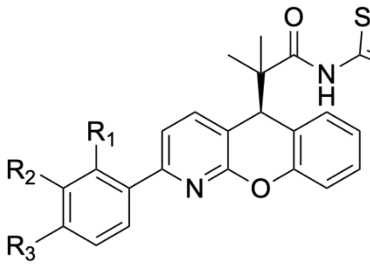


| Compounds | Substitution Site | Functional Group |
|---|---|---|
| **14** | - | - |
| **15** | R1 | Me |
| **16** | R2 | Me |
| **17** | R3 | Me |
| **19** | R3 | OMe |
| **20** | R3 | Et |
| **21** | R3 | *n*Pr |
| **22** | R3 | O$^{i}$Pr |
| **23** | R3 | OH |
| **24** | R3 | $NMe_2$ |
| **26** | R3 | Ac |
| **27** | R2 | F |
| | R3 | O$^{i}$Pr |
| **28** | R2 | F |
| | R3 | Ac |

**Table S8.** Chemical structures of TNI-97 compound series used for AFES calculations.

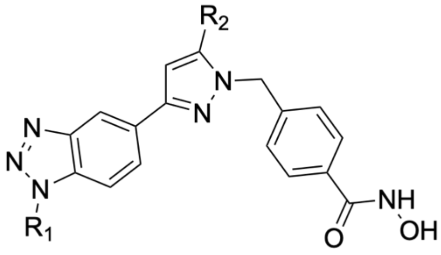


| Compounds | R1 | R2 |
|---|---|---|
| **30a** | $-CH_3$ | $CF_3$ |
| **30b** | | Ph |
| **30c** | | 4-$CF_3$Ph |
| **30d** | | 2-$CH_3$Ph |
| **30e** | | 3-$CH_3$Ph |
| **30f** | | 4-$CH_3$Ph |
| **30g** | $-C_2C_5$ | $CF_3$ |
| **30h** | $-C_2C_5OCH_3$ | $CF_3$ |
| **30i** | $-CH(CH_2)_2$ | $CF_3$ |
| **30j** | | Ph |
| **30k** | | 4-F-Ph |
| **30l** | | 4-$CF_3$Ph |
| **30m** | | Bn |
| **30n** | -Bn | $CF_3$ |
| **30o** | | 4-$CF_3$Ph |
| **30p** | | 2-$CH_3$Ph |
| **30q** | | 3-$CH_3$Ph |
| **30r** | | 4-$CH_3$Ph |

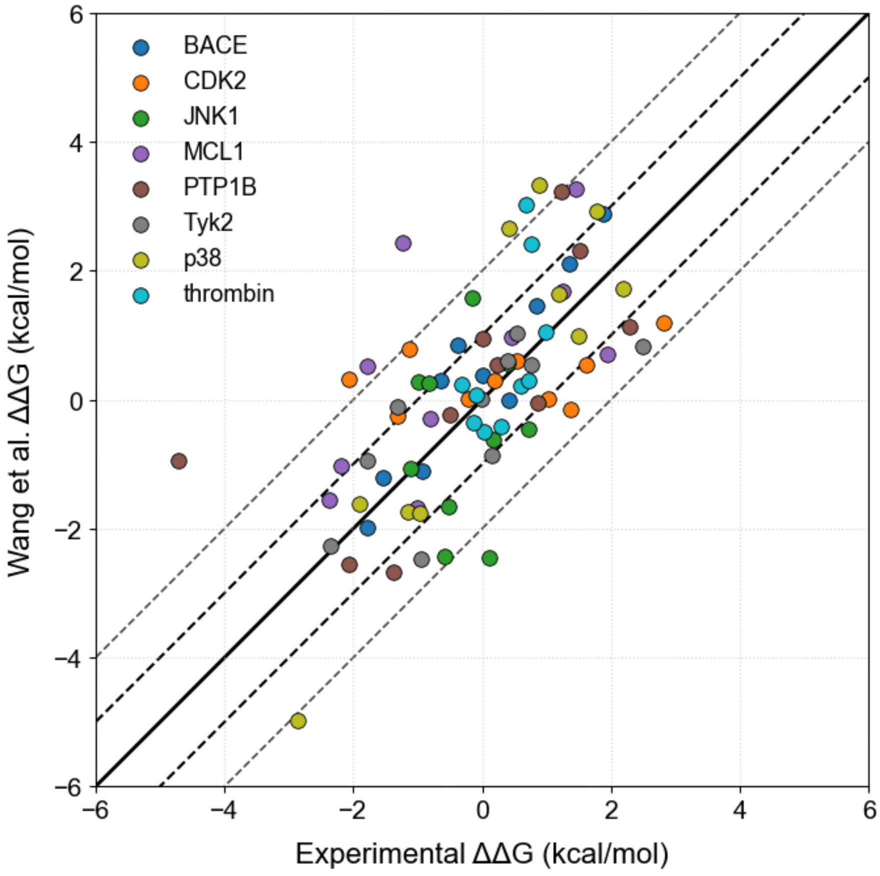


**Figure S1.** $\Delta\Delta G_{bind}$ prediction-experiment reported by Wang et al..

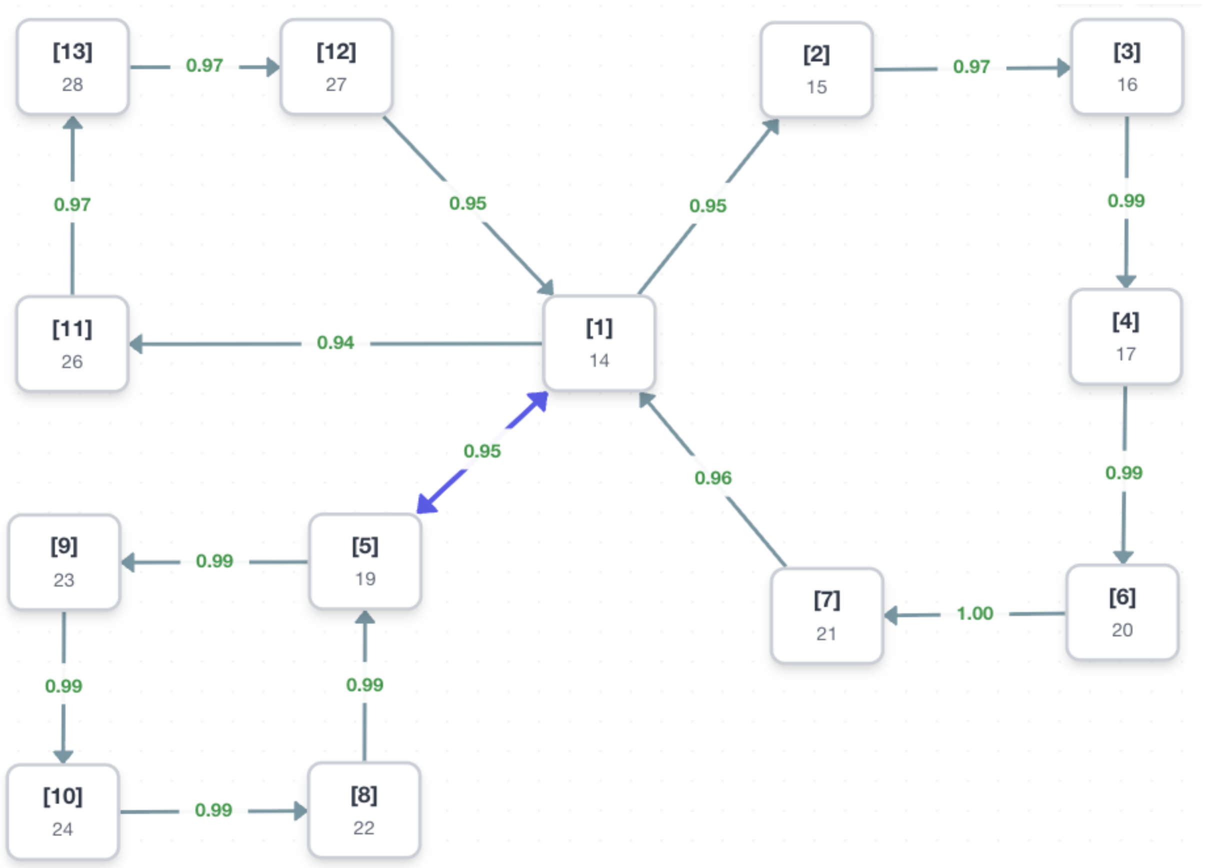


**Figure S2.** Ligand perturbation map for the glucocorticoid receptor (3BQD) system. Each node represents a ligand, labeled with its compound name (in **Table S7**), and arrows indicate the direction of the relative free energy perturbations. The numerical value displayed near each arrow represents the similarity score between the connected nodes. Multiple closed cycles were generated by connecting ligands with similar functional groups or heteroatoms to ensure high phase-space overlap. The compound numbering starts from [1] on the map corresponding to compound 14.

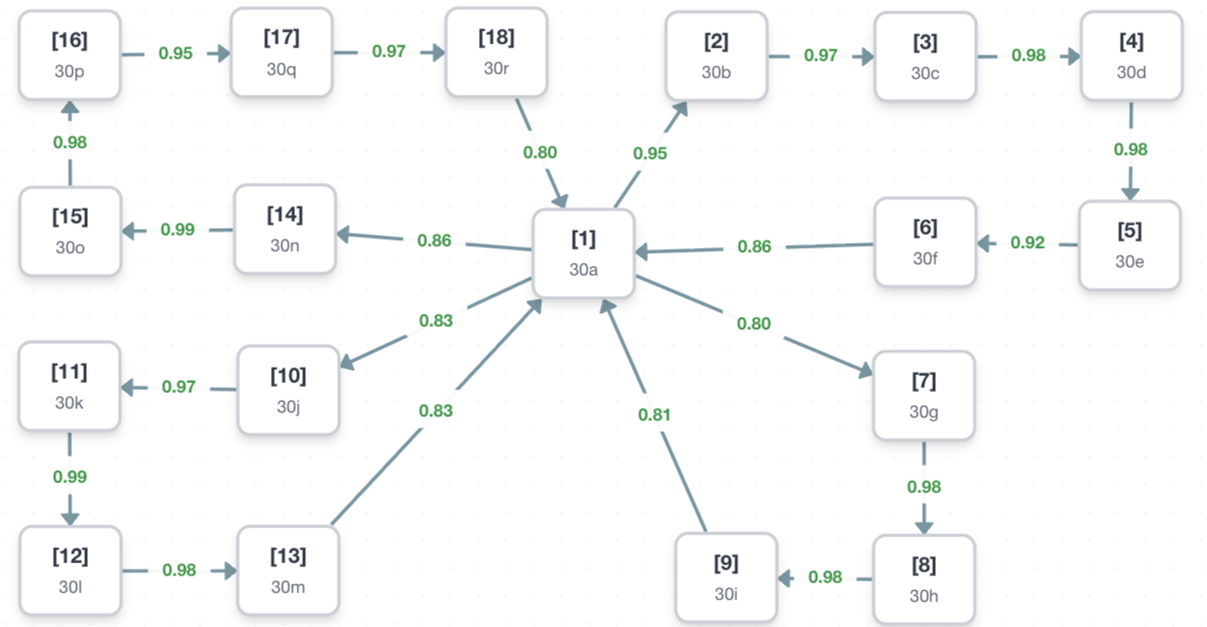


**Figure S3.** Ligand perturbation map for the HDAC6 inhibitor (5EDU) system. Each node represents a ligand, labeled with its compound name (in **Table S8**), and arrows indicate the direction of the relative free energy perturbation. The numerical value displayed near each arrow represents the similarity score between the connected nodes. Multiple closed cycles were constructed between ligands bearing diverse R1 and R2 substituents to ensure high phase-space overlap. The compound numbering starts from [1] on the map corresponding to compound 30a.
